# Benchmarking RNA velocity methods across 17 independent studies

**DOI:** 10.1101/2025.08.02.668272

**Authors:** Ya Luo, Jun Ren, Qian Yang, Ying Zhou, Zhiyu You, Qiyuan Li

## Abstract

RNA velocity techniques offer great potential for unveiling trajectories of cell state transitions in different biological contexts. While diverse computational methods have been developed, there is no evidence-based guidelines for best-practice in RNA velocity inference. Here we conducted a benchmark study for 14 existing RNA velocity methods in 17 independent datasets. Many validations were done for the first time. We evaluated the performances of each method by measuring accuracy, stability, and usability. Our data showed no single method exhibited superior performance in all the assessments, and unexpected underperformance was observed in certain cases. Especially, the lack of uniformity in the inference results highlights the necessity to compare and control of multiple methods in a single analysis. Our study revealed current limitations and challenges in the RNA velocity methods and informed the best-practice for future studies.

## Introduction

Single-cell RNA sequencing (scRNA-seq) has enabled a high-resolution snapshot of the cellular status and transitions, facilitating the development of computational approaches unveiling the transcriptional dynamics in diverse biological contexts^1-3^. RNA velocity has become a powerful technique in investigating cell fate determination by offering comprehensive views of the trajectories of cells during state transitions.

RNA velocity recovers information about cellular dynamics by measuring the abundance of unspliced and spliced mRNA at single-cell level^4, 5^, which can be explicitly quantified using various established tools, such as velocyto^4^, STARsolo^6^, and dropEst^7^. RNA velocity inference generally comprises three stages: pre-processing, velocity estimation and post-processing (Fig.1A). In the pre-processing stage, spliced and f unspliced mRNA abundance matrices are initially filtered to keep only highly variable genes, which are subsequently normalized, projected to lower dimension and smoothed. The velocity estimation steps vary substantially among different methods. Many methods infer RNA velocity based on the steady-state assumption, as implemented in velocyto^4^. The other methods infer RNA velocity by regressing spliced and unspliced mRNA abundance to the trajectories defined by optimized dynamic models, as seen in scvelo^5^. The steady-state model, such as velocyto, assumes: 1) gene transcription and degradation rates are constant, and splicing rates are uniform for all genes; 2) a subset of cells reside in steady-state. Both assumptions are circumstanced to large, homogeneous cell populations. Then, the method such as scvelo offers a dynamical model based on maximum likelihood estimation that relaxes the steady-state assumption by introducing cell specific latency time for each gene, and the kinetic parameters are jointly estimated to enhance the flexibility and accuracy. In the post-processing stage, gene-wise RNA velocity vectors are projected into a low-dimensional space using Uniform Manifold Approximation and Projection (UMAP), t-distributed stochastic neighborhood embedding (t-SNE), or principal component analysis (PCA), to ultimately demonstrate the trajectory map.

Up to date, more than 20 RNA velocity methods are published, which explicably adopted either of the following methodologies (Fig. 1B). The first is to improve the accuracy of inference by incorporating additional biological information. For examples, protaccel^8^ integrates protein translation processes into the model; Chromatin Velocity^9^ and MultiVelo^10^ incorporate epigenomic features like chromatin accessibility; PhyloVelo^11^ combines genealogical information with single-cell transcriptome data for velocity field reconstruction; Dynamo^12^ and velvet^13^ utilize time-resolved single-cell metabolic marker data to model dynamics gene expression. The second methodology focuses on improving the algorithms to better control the noises and specifically recapitulate the dynamics of the transcription process. These include veloVAE^14^, veloVI^15^ and Pyro-Velocity^16^ that adopt Bayesian inference framework to cope with uncertainty. Then, veloAE^17^ smooths low-dimensional velocities with an autoencoder; Deepvelo^18^ that applies a variational autoencoder to predict continuous changes in intracellular gene expression; uniTVelo^19^ develops a uniform regularization of velocity estimation to control for biases in time scale and directionality; Deepvelo^20^ utilizes graph convolutional networks to infer gene-specific and cell-specific kinetic parameters; latentvelo^21^ constructs a neural ODE model in the presence of a multi-branching temporal process; cellDancer^22^ uses deep neural network to predict cell-specific kinetic parameters for each gene, thereby enabling accurate RNA velocity estimation across different cell types; finally, cell2fate^23^ decomposes the velocity matrix into resolvable modules representing the biological factors underlying transcriptional dynamics.

**Fig. 1:**
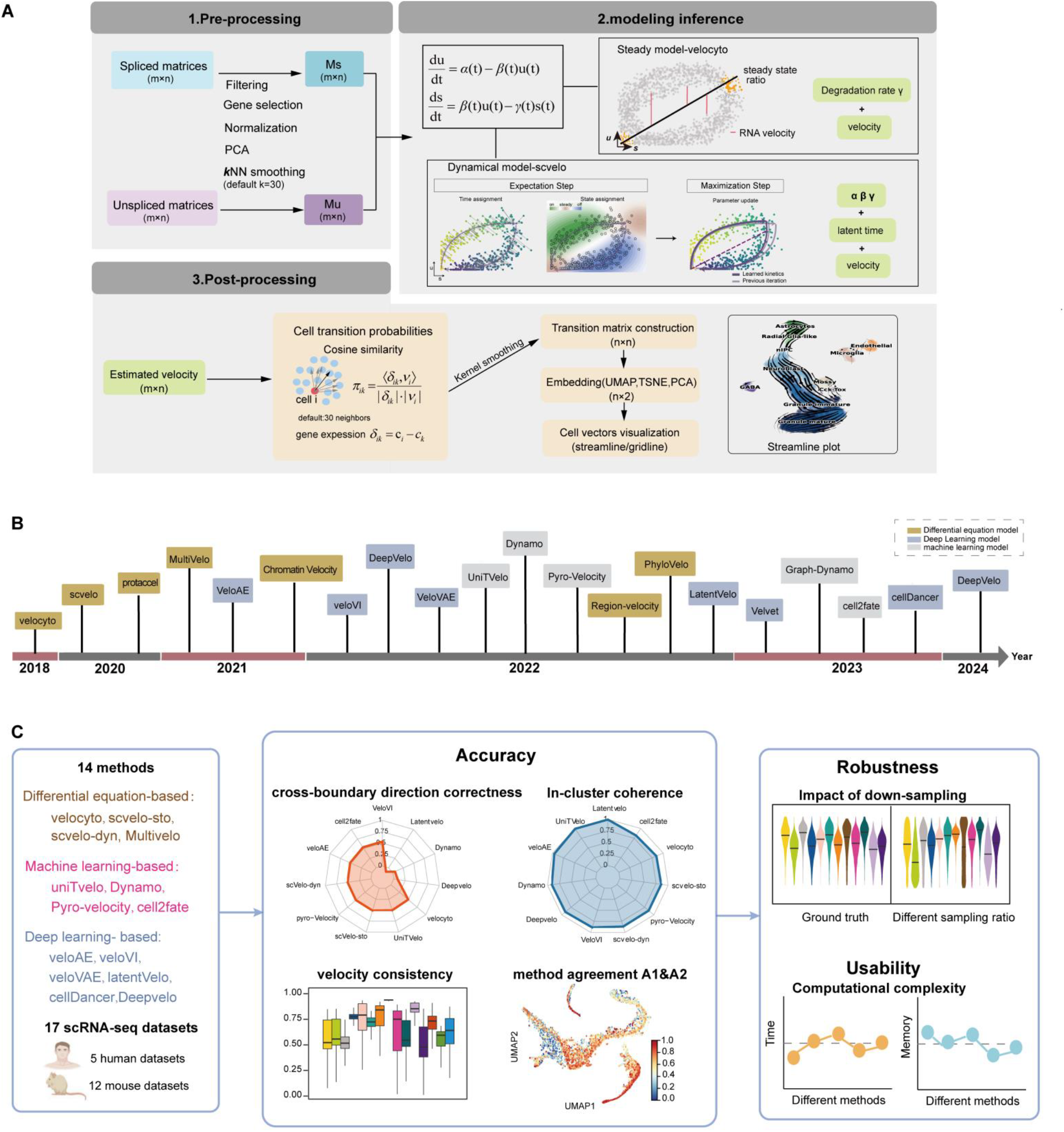
An overview of RNA velocity methods. **(A)** Summary of three-step RNA velocity analysis. **(B)** Timeline of RNA velocity methods. **(C)** Schematic plot of the benchmarking workflow for 14 RNA velocity methods in the 17 scRNA-seq datasets.

The rapid advancement of RNA velocity methods has provided researchers with diverse analytical tools, yet it also leads to the uncertainty and difficulty in choosing the best-practice for specific analyses. Previous studies^24-26^ compared the prevalent RNA velocity methods, but the evaluation was limited by the number of methods and datasets. Here, we conducted a systematic evaluation of 14 RNA velocity methods, assessing the performance in full-scale including inference accuracy, algorithmic stability, and computational resource usage. Our benchmark study utilized 17 independent datasets, in which most methods were tested for the first time (Table 1, Methods). By rigorously comparison of the performance across multiple datasets and conditions, our data offer insights for the current RNA velocity methods and inform the best-practice for future studies.

**Table 1.** Overview of cross-species single-cell sequencing datasets for benchmarking test.

| Original Identifier | Species | Technology | Description | Number of cells | Number of genes | Dataset No. |
| --- | --- | --- | --- | --- | --- | --- |
| GSE132188 | Mus musculus | 10x Genomics | Pancreatic endocrinogenesis | 3696 | 27998 | Dataset1 |
| GSE95753 | Mus musculus | 10x Genomics | Dentate gyrus | 2930 | 13913 | Dataset2 |
| GSE87038 | Mus musculus | 10x Genomics | Mouse Erythroid Maturation | 9815 | 53801 | Dataset3 |
| ERP120467 | Homo sapiens | 10x Genomics | Human Bone Marrow | 5780 | 14319 | Dataset4 |
| GSE128365 | Mus musculus | scEU-seq | Intestinal organoid | 3831 | 9157 | Dataset5 |
| GSE122466 | Mus musculus | 10x Genomics | Mouse retina development | 2726 | 31053 | Dataset6 |
| GSE118068 | Mus musculus | 10x Genomics | Mouse hindbrain (GABA,Glial) | 13501 | 55421 | Dataset7 |
| GSE119945 | Mus musculus | 10x Genomics | Mouse organogenesis (chondrocyte) | 30000 | 55401 | Dataset8 |
| GSE162170 | Homo sapiens | 10x Genomics | Developing human brain | 4693 | 954 | Dataset9 |
| SRP129388 | Homo sapiens | 10x Genomics | Human postconception week 10 forebrain | 1720 | 32738 | Dataset10 |
| SRP073767 | Homo sapiens | 10x Genomics | PBMC-68k | 65877 | 33939 | Dataset11 |
| GSE209878 | Homo sapiens | snRNA-seq | Human HSPC | 11605 | 1000 | Dataset12 |
| GSE141851 | Mus musculus | scNT-seq | Mouse cortical neuronal | 3066 | 44021 | Dataset13 |
| SRP135960 | Mus musculus | 10x Genomics | Mouse hindbrain (Oligodendrocyte) | 6253 | 27998 | Dataset14 |
| GSE109989 | Mus musculus | inDrop | Mouse bone marrow | 2600 | 21552 | Dataset15 |
| www.10xgenomics.com | Mus musculus | 10x Genomics | Embryonic E18 Mouse Brain (5k) | 3653 | 32285 | Dataset16 |
| GSE81682 | Mus musculus | 10x Genomics | Mouse hematopoiesis | 1772 | 3970 | Dataset17 |

## Results

### Design of the benchmark study

To fully represent the state-of-the-art in RNA velocity methods, our benchmark study included 14 algorithms that are publicly available (Table 2, Methods). These methods are categorized into three groups: 1) ODE-based methods including velocyto^4^, scvelo-stochastic^5^, scvelo-dynamic^5^, and Multivelo^10^; 2) machine learning-based methods such as uniTvelo^19^, Dynamo^12^, Pyro-velocity^16^, and cell2fate^23^; and 3) deep learning-based methods like veloAE^17^, veloVI^15^, veloVAE^14^, latentVelo^21^, cellDancer^22^, Deepvelo^20^ (Fig.1C)

**Table 2.**
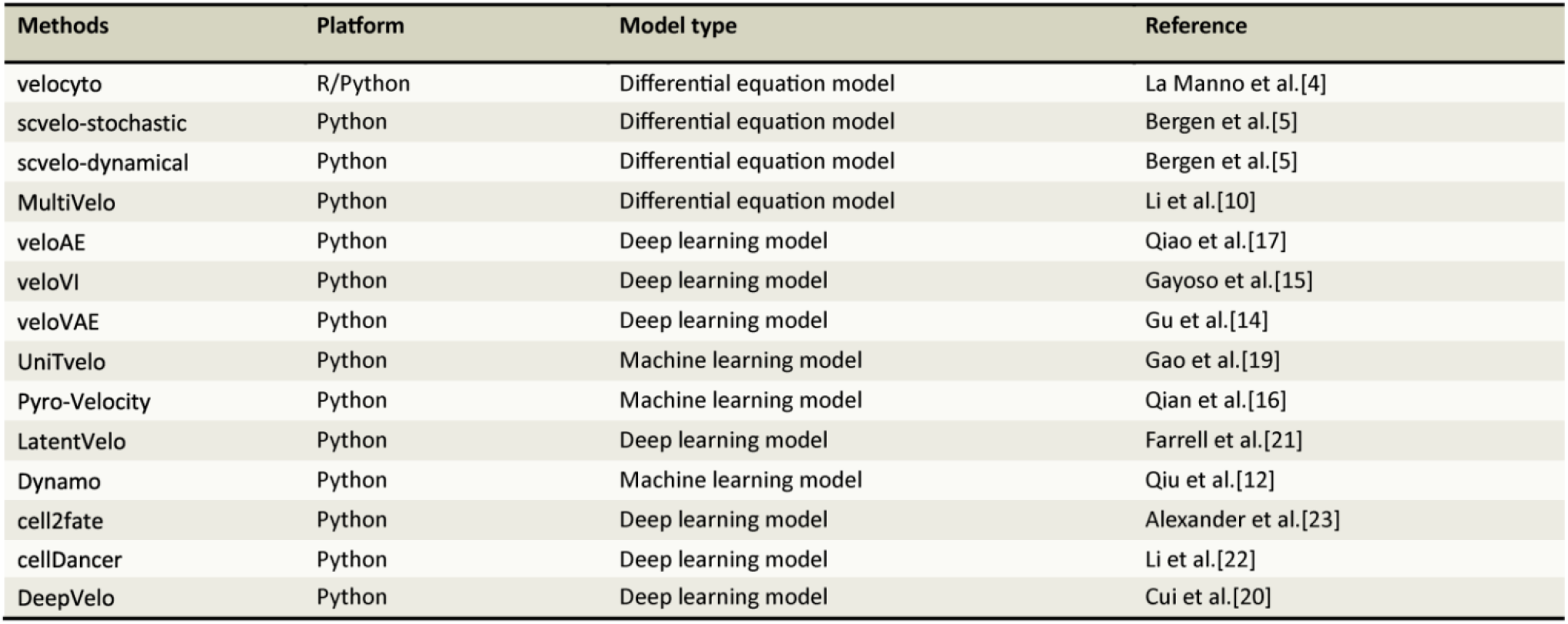
Description of the 14 RNA-velocity methods.

As for the datasets, we collected 17 published single-cell RNA sequencing (scRNA-seq) datasets, covering a diverse biological contexts and sequencing technologies (Table 1, Methods). Particularly, certain datasets (dataset 1-5 and dataset 10) were extensively utilized for evaluating the performance of RNA velocity methods, while other datasets (dataset 9, dataset 12 and dataset 17) were tested only once (Supplementary Fig.1). For performance evaluation, we used four metrics. Cross-boundary direction correctness (CBDir)^17^, intra-cluster coherence (ICCoh)^17^, and velocity consistency^5^ measure the correctness and coherence of inference at cell level by each method. Then, method agreement A1 & A2^25^ measure the consistency of inference among different methods. We systematically benchmarked the performance of 14 RNA velocity methods in all 17 datasets. In addition, we conducted down-sampling experiments on four representative datasets to assess the stability of the methods. Lastly, we conducted a comparative analysis of the computational resource consumption of each method in practical applications, including time and memory requirements (Fig. 1C).

### The performance of RNA velocity inference varies substantially among different methods

We compare the performance of 14 methods on velocity estimation in 17 independent datasets. The implementation of each algorithm followed strictly the description of the original study. (Methods). We examined the results of each method measured on different datasets. As a result, the test performance of the RNA velocity methods showed substantial variation in all aspects.

For CBDir, with an average of 0.14, the test results suggested that for most of the RNA velocity methos, there is still a large room for improvement. veloVI demonstrated the highest overall performance (CBDir=0.28), followed by pyro-velocity (CBDir=0.24) (Fig.2A). Nevertheless, we noticed that most of the inferred transitions by veloVAE were reversed, as evidenced by negative CBDir values (Fig.2A, Supplementary Fig.2). Besides, the consistency and coherence for RNA-velocity prediction tended to decline with the complexity. In the human bone marrow cells dataset (i.e., dataset4) with multiple transcriptionally enhanced cell trajectories, the average CBDir was minimized to -0.154 (Supplementary Fig.3A). In the mature-state peripheral blood mononuclear cell dataset (i.e., dataset11), most methods yielded a number of erroneous directions, such as the transition from CD8+Gytotoxic T cells to CD4+/CD45RA+/CD25-Naive T cells, which is contrary to the known biology^27^ (Supplementary Fig.3A, 4). In addition, some of the methods yielded higher CBDir values in datasets included in the original study than in datasets where they were tested for the first time. (Supplementary Fig.5A).

**Fig. 2:**
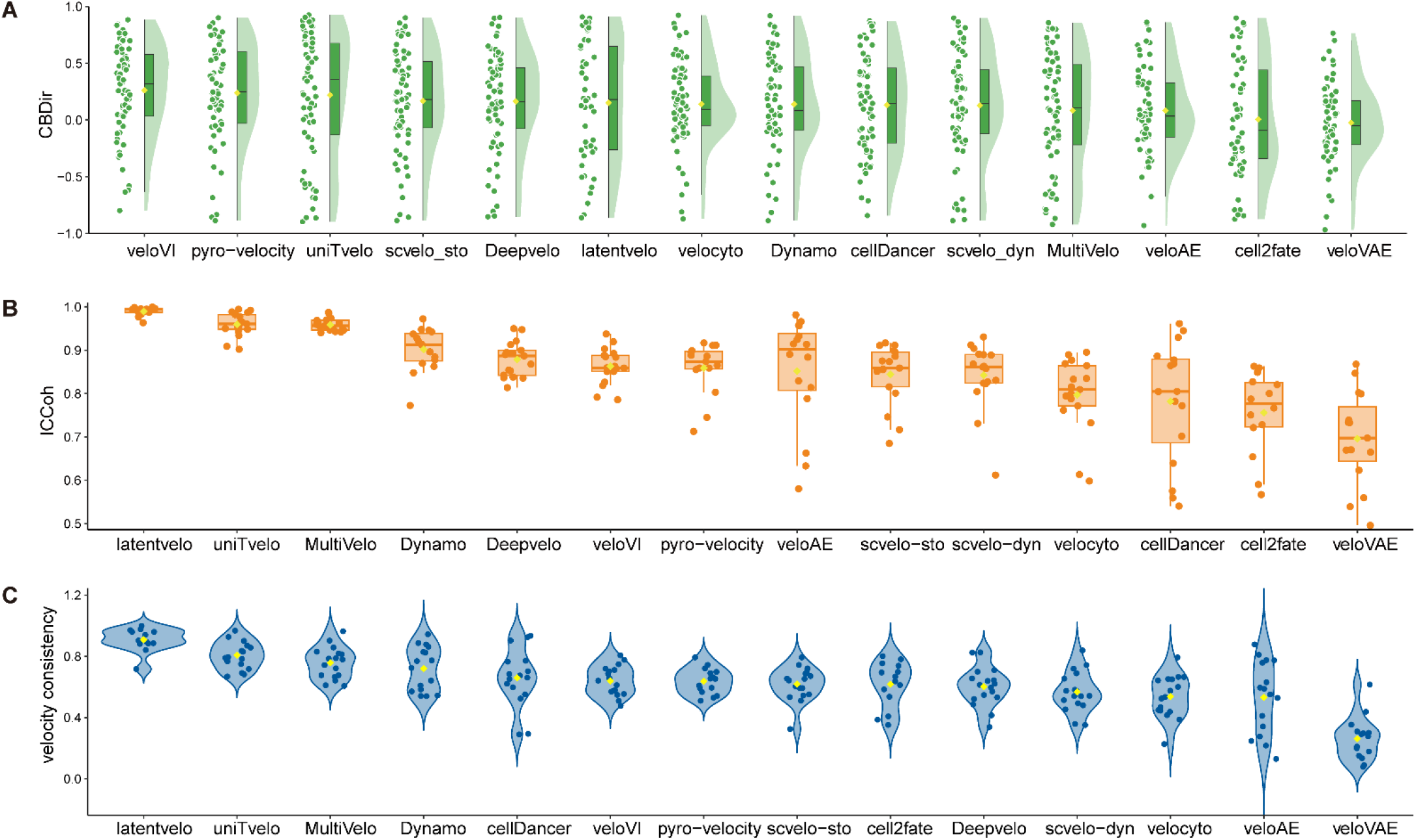
The performance of 14 RNA velocity methods across 17 scRNA-seq datasets. **(A)** Cross-boundary Direction Correctness (CBDir) scores. Each point represents the directional correctness score for a source-to-target cluster transition (e.g., cluster A → cluster B) within a dataset. **(B)** Intra cluster coherence (ICCoh) scores. **(C)** Velocity consistency scores. Individual points represent the respective metric’s mean score within a single dataset (For B & C). Yellow symbols denote the grand mean of each method’s scores for the corresponding metric, calculated across all datasets.

On the other hand, most methods achieved relatively high ICCoh values (>=0.7), especially latentvelo (ICCoh=0.99), uniTvelo (ICCoh= 0.96), and MultiTvelo (ICCoh= 0.96). Likewise, these methods also demonstrated high velocity consistency (>=0.6). Both metrics indicated that the inferred velocity fields were quite smooth among neighboring cells (Fig.2B, C). Nevertheless, veloVAE underperformed in both intra-cluster coherence and velocity consistency compared to other methods (Fig.2B, C, Supplementary Fig.3B, C). Moreover, for all the methods, the mean ICCoh showed no biases for different test datasets (Supplementary Fig.5B). Whereas for velocity consistency, most methods performed significantly better in the datasets used in the original studies (Supplementary Fig.5C). To summarize, our data suggested that existing methods infer consistent velocities at single-cell level, but the results demonstrate substantial inadequacy in resolving the transitions among different cell states hence limited ability to distinguish truly relevant transcriptional dynamics.

### Discrepancies in the velocity fields inferred by different methods

Then, we evaluated the consistency in the velocity fields inferred by 12 different methods using two metrics of agreement, A1 and A2^25^ (Dynamo and cellDancer were excluded for the related functionality was unsupported, Methods).

Our data showed that the agreement scores (A1) of the 12 methods were generally low (A1<0.3), indicating substantial discrepancies in RNA velocity estimations among these methods. In particular, latentvelo and cell2fate manifested quite low consistency scores A1 to all the other methods and between each other (Fig.3A, Supplementary Fig.6,7). We further investigated method agreement scores within different cell types for four methods with relatively high consistency (velocyto, scvelo-dyn, veloVAE, and Deepvelo, Methods). In four widely used benchmark datasets (dataset1-4), we found that the inferred RNA velocity by these methods were more consistent for cells in the early-stage of differentiation, such as ductal cells, ngn3 low or high endocrine progenitors and endocrine precursors. While for highly differentiated cell types (Epsilon cells, Alpha cells, Beta cells), the inference consistency dropped about 64% on average among the four methods (Fig. 3B, C, Supplementary Fig.8A). In the human bone marrow dataset, only veloVI and Deepvelo exhibited consistency across all cell types (Supplementary Fig.8B), while the consistency of the methods across cell types was generally lower in the other two datasets (Supplementary Fig.8C, D).

**Fig. 3:**
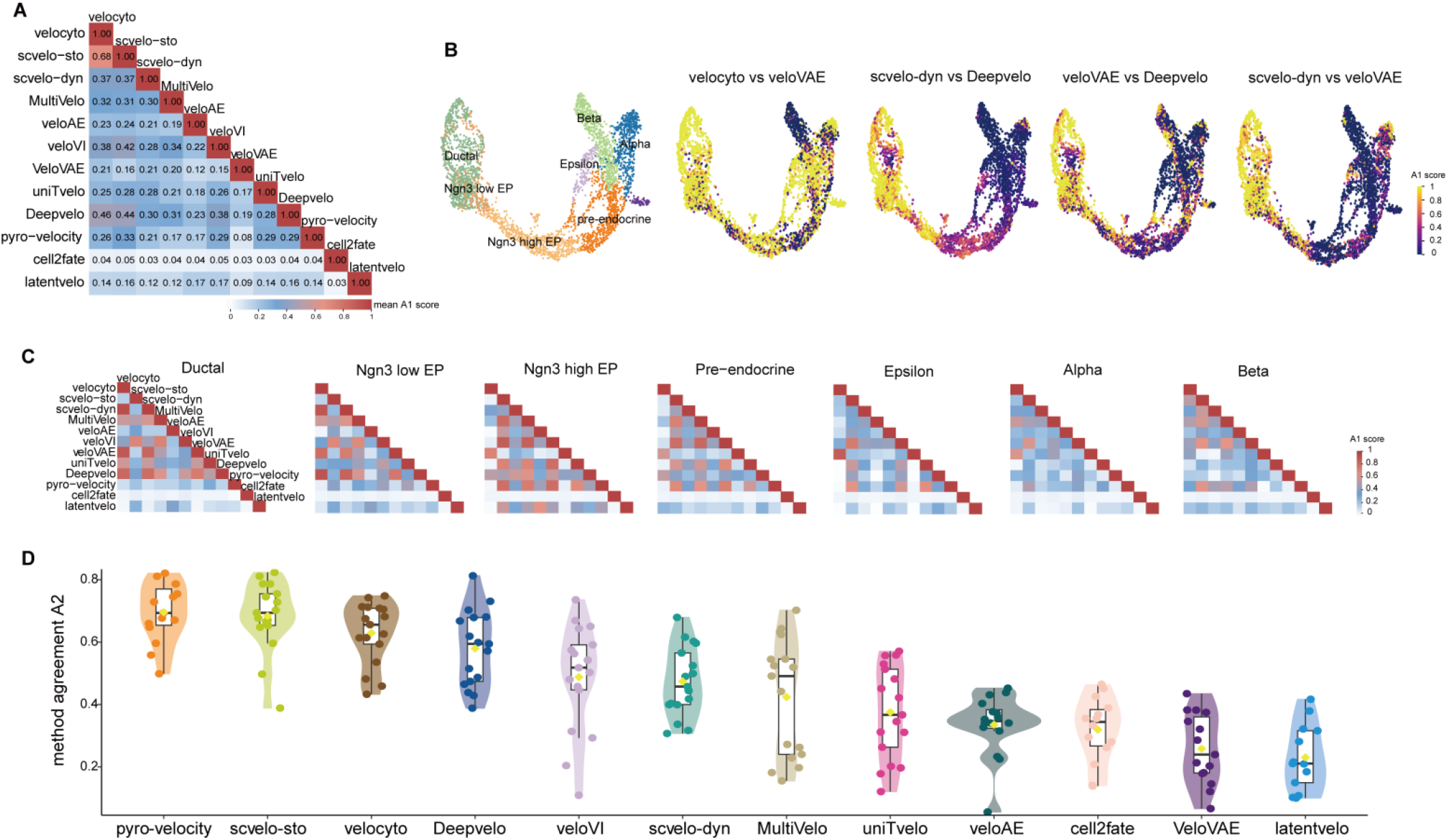
Comparison of cell velocity field agreement for RNA velocity methods. **(A)** Pairwise comparison of method agreement A1 mean scores. **(B)** Dataset 1 (pancreatic endocrinogenesis): UMAP embedding plots colored by cell type (left) and method consistency A1 score for selected pairwise methods (right). **(C)** Pairwise comparisons of method consistency A1 scores for the seven cell types in dataset1 (pancreatic endocrinogenesis). **(D)** Method agreement A2 mean scores of 12 RNA velocity methods across all datasets. Individual data points (circles) represent the mean A2 score per dataset for each method. Grand means (±SD) across datasets are indicated by yellow symbols.

Next, we assessed the systematic differences (A2) among these methods. As a result, four methods (pyro-velocity, scvelo-sto, velocyto, and Deepvelo) performed relatively better (A2>0.5) than others, and latentvelo, again, exhibited lowest consistency (A2<0.3) (Fig.3D, Supplementary Fig.9). In the four widely used benchmark datasets, scvelo-sto and pyro-velocity demonstrated higher consistency (A2>0.7), while cell2fate and latentvelo were the least consistent (A2<0.35) (Supplementary Fig.10-11).

In summary, our data suggested that discrepancies in RNA velocity estimation are quite common among the tested methods, especially in certain methods and cell types. Consequently, to mitigate potential bias introduced by a single method, it is advisable to integrate results from multiple methods in studies related to RNA velocity.

### Stability of RNA velocity inference is subject to sampling biases

Current methods infer RNA velocity based on measures of RNA abundance from large number of genes and cells hence the results are subject to all technical and sampling biases. Here, we assessed the stability of the existing methods by inferring RNA velocity from various down-sampling cell populations (Dataset 1-4).

We evaluated the stability of the performance of each method in the down-sampling of cells based the same metrics above. In general, CBDir is the metric that was affected the most by varying sampling rates. For most of the methods, CBDir varied drastically at different sampling rates and the tendencies of the change were largely unpredictable (Fig. 4A, Supplementary Fig.12A, C). Specifically, our data suggested that pyro-velocity and cell2fate were the most unstable for CBDir (range [-0.11, 0.403]). On the other hand, ICCoh were relatively stable at different down sample rates (above 0.7), except for some outliers (velocyto and cellDancer, Fig.4B, Supplementary Fig.12B, D). As for the velocity consistency, the performance of the methods tended to polarize in the down-sampling. Methods such as uniTvelo and latentvelo remained stable and performed well at different sampling rates, while other methods tended to be unstable and underperformed (Fig. 4C, Supplementary Fig.13).

**Fig. 4:**
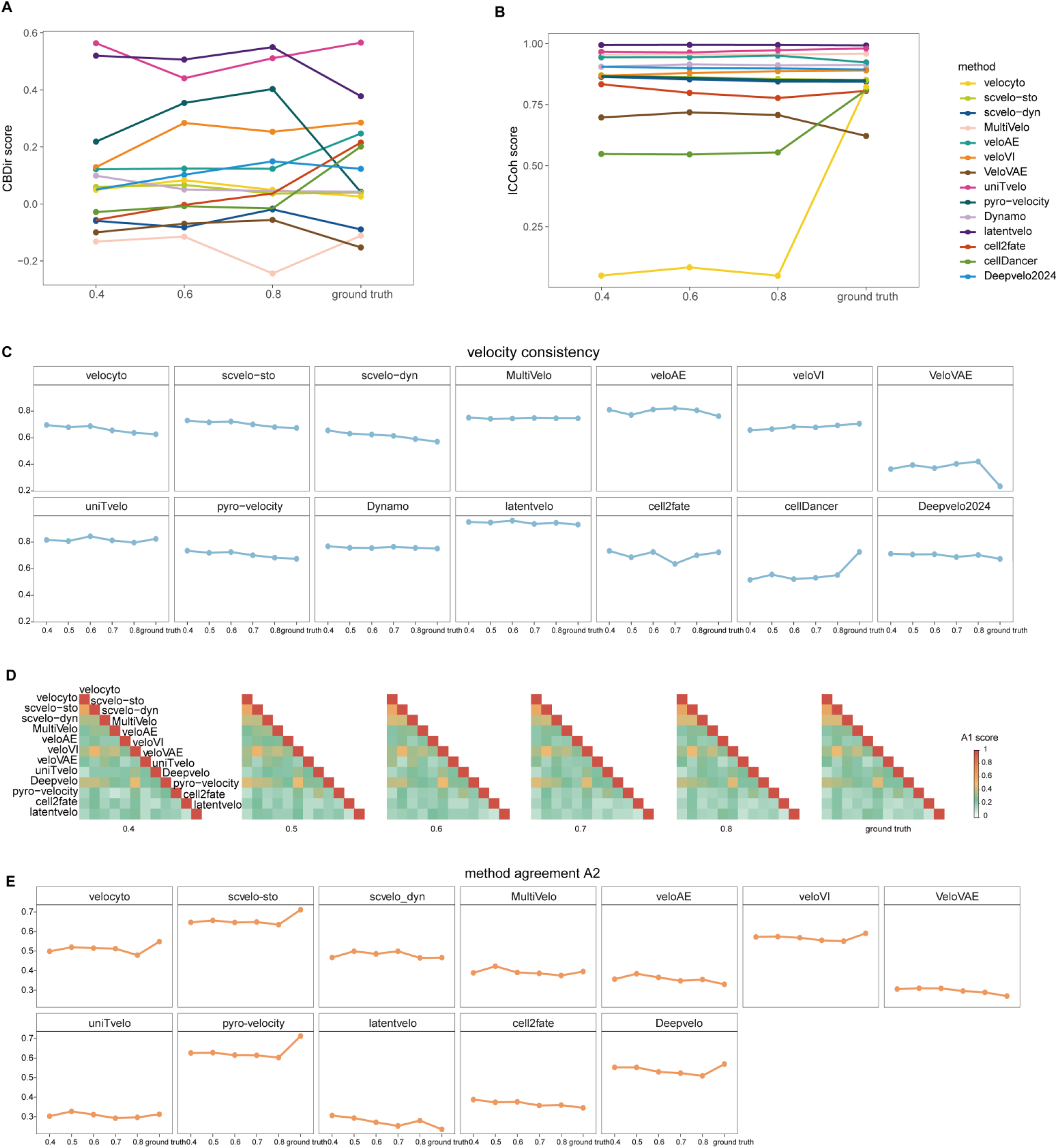
Comparing the stability of RNA velocity methods for down-sampling from the original four datasets. **(A)** The average of CBDir score **(B)** ICCoh score **(C)** and velocity confidence score of four benchmark datasets were processed by 14 RNA velocity methods with different sampling rate. **(D)** The average of the method agreement A1 score of the four benchmark datasets processed by the 12 RNA velocity methods with different sampling rate. **(E)** and the average of method agreement A2 score. Ground truth represents the original dataset.

Regarding the inter-method agreement A1, despite of different sampling rates, the overall agreement of inferred RNA velocity by these methods remains unchanged. Notably, some methods, including Deepvelo, scvelo-sto, veloVI, and velocyto, showed highly stable agreement (Fig. 4D). As for the agreement A2, most of the methods exhibited stable inter-method agreement, drop only slightly in the down-sampling data. Nevertheless, methods achieved high agreement in full data also tend to agree in the down-samples, such as scvelo-sto and pyro-velocity (A2>0.5) (Fig. 4E, Supplementary Fig.14).

### Overall performance

Next, we derived comprehensive scores for the methods in three aspects of performance based on summary of the metrics, 1) the accuracy of RNA velocity estimation (CBDir, ICCoh and velocity consistency, and inter-method agreement A2); 2) the stability of inference, based on performance in the down-sampling data; and 3) the usability of all methods, namely, the computational efficiency.

Our scores revealed that none of the methods emerged as the top performer in all aspects. As standalone methods, veloVI outperformed the other methods in terms of inference accuracy (Fig. 5A), and latentvelo turn out to be the most stable performer at various sampling rates (Fig. 5B). In terms of inter-method agreement, pyro-velocity and scvelo-sto showed high agreement with current methods, and also turned out to be the most stable (Fig. 5A, B).

**Fig. 5:**
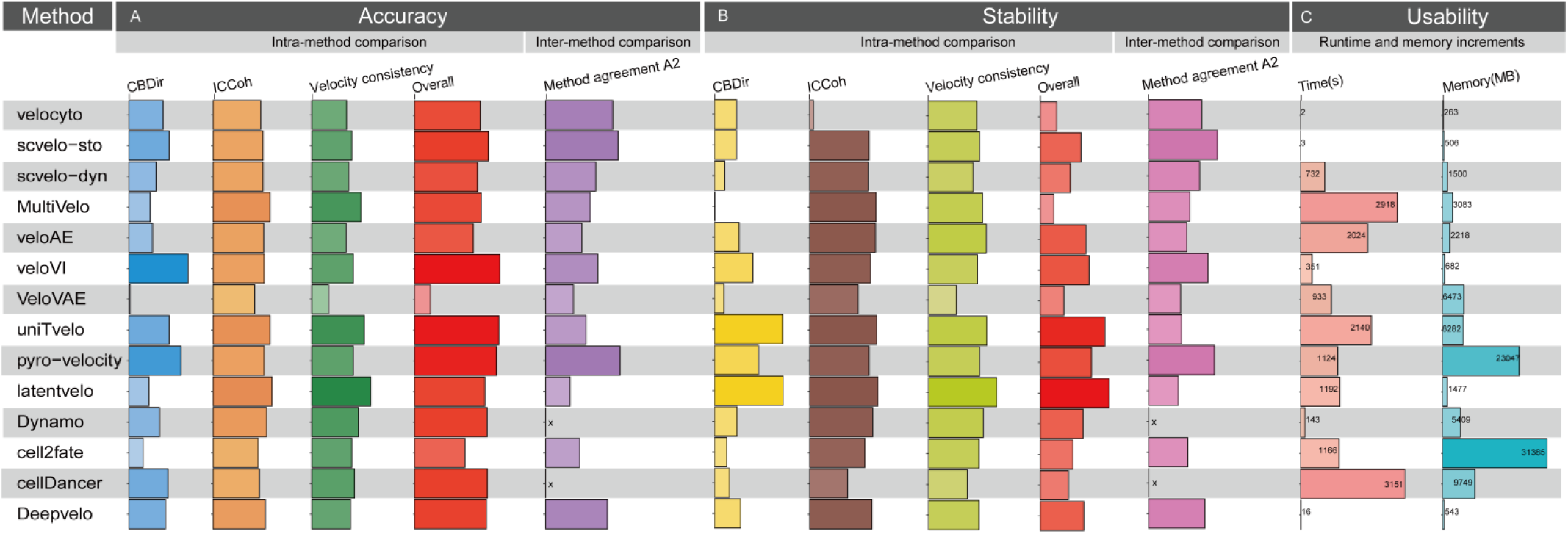
Summary of performance of all methods. The method information includes intra-method evaluation metrics (cross-boundary direction correctness, intra-cluster coherence and velocity consistency) as well as inter-method evaluation metrics (method agreement A2); stability evaluation (cross-boundary direction correctness, intra-cluster coherence and velocity consistency and method agreement A2); and runtime and memory increments. Where ‘overall’ calculates the geometric mean score of three metrics (cross boundary direction correctness, intra-cluster coherence and velocity consistency).

As for computational efficiency, Deepvelo stood out as the most efficient RNA velocity method, with an average GPU computation time of 16 seconds and an average memory increment of only 543MB, followed by veloVI and Dynamo. Notably, pyro-velocity and cell2fate consumed a large amount of GPU memory, with an average memory increment exceeding 22GB (Fig. 5C).

## Discussion

RNA-velocity analysis is gaining traction in better resolving the complex dynamics of cell-state transitions implicated in diverse biological contexts. In this benchmarking study, we tested 14 RNA velocity methods in 17 published datasets, conducting a comprehensive performance evaluation in the dimensions of inference accuracy, algorithm stability, and computational efficiency. Our results suggested that the performance of RNA velocity methods is significantly influenced by differences in cell characteristics. As our data showed that with increasing complexity of transcriptional dynamics, the inference performance of the methods tended to decline. For example, in a peripheral blood mononuclear cell dataset at a mature state, many methods inferred incorrect directions; in a human bone marrow cell dataset, due to the presence of multiple transcriptionally enhanced cell trajectories, most methods exhibited low cross-boundary direction correctness (CBDir). Additionally, cell lineage and differentiation state also influence the performance of RNA velocity methods. For example, in the pancreatic endocrinogenesis dataset, early progenitor cells with gradual transcriptional changes were accurately resolved by most of the methods, whereas consistency among the methods decreased drastically in the same population of cells but at highly differentiated levels, highlighting the challenges posed by the complexity of differentiation lineages for velocity inference.

Nevertheless, the current benchmarking analysis has some limitations. On the one hand, due to the fast advances in the field, some new methods were not included in our benchmarking study. On the other hand, while our metrics provide multi-dimensional insights of the inference performance, there are some key trade-offs. First, CBDir is used to measure the accuracy of cell directional transitions. However, CBDir relies on established ground truth, yet for most single-cell datasets, CBDir was computed based on curated trajectories of cells, which is subject to errors or insufficient knowledge. Second, we used ICCoh and velocity consistency to measure the coherence of inferences, for which most methods achieving high scores (ICCoh>=0.7, and velocity consistency>=0.6). However, such high scores can be resulted from over-smoothing of the trajectory rather than real biological continuity and introduce bias against bifurcative and multifurcative trajectories. Then, the method consistency metrics (A1 & A2) assesses the consistency of inferred velocity fields across different methods. Our data showed that discrepancy is very common in the RNA velocity inferred by different methods, particularly for certain cell types. However, such discrepancy can reflect different biology context embedded in the different algorithms. For example, latentvelo and cell2fate exhibit significantly discrepancy, which is attributed to the unique model architecture.

Furthermore, we performed simple down-sampling (ratio) of the raw data to simulate sampling biases and insufficient observations, but the actual biases in real data are more complex. Nevertheless, our data showed that sampling biases, such as insufficient sampling of rare cell types or intermediate transitional cells, lead to erroneous or incomplete trajectory reconstruction. Data sparsity is a fundamental limitation of traditional scRNA-seq technology. Integrating metabolic labeling single-cell sequencing technology^28-30^ or other multi-omics data (e.g., chromatin accessibility or protein translation) will improve the quantification of spliced and un-spliced mRNA. Again, integration of other data in RNA velocity study is constrained by low throughput, complex experimental design, and high costs.

In summary, our benchmarking results informed future directions to improve RNA velocity methods. First, large, well-balanced, multi-center datasets by combining lineage tracing or metabolite labeling techniques will mitigate technical biases and provide foundational benchmarking data for validation of the methods. Second, parallel benchmarking of the methods is needed to suggest best-practice in RNA velocity inference. Third, better control of the imbalance in the cell populations will improve the inference performance. Finally, establishing a consensus method framework to integrate predictions from multiple RNA velocity methods is highly recommend for control of biases introduced by any single method, thereby providing more reliable velocity inference.

## Methods

### Benchmark metrics

We established a common pipeline to comprehensively assess the performance of RNA velocity methods across the 17 datasets. In the pipeline, we used the following four metrics to evaluate each method.

#### 1. Cross-boundary direction correctness (CBDir) ^17^

CBDir is used to assess the consistency of the velocity direction of a cell with the expected direction of development within a brief timeframe. Specifically, given the ground truth development direction e.g. A → B, the correctness of the transition from the source cluster to the target cluster is measured by considering the boundary cells that reflect the development of the cell over a short period of time. The boundary cells represent the set as

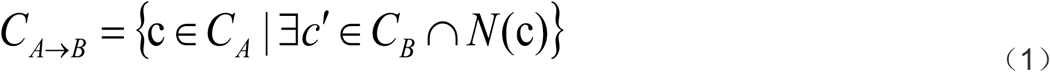

where C_A_ is sets of cells in source cluster A and C_B_ is sets of cells in target cluster B, *N*(*c*) stands for the neighboring cells of specified cell *c*.

The CBDir value was calculated using the following equation:

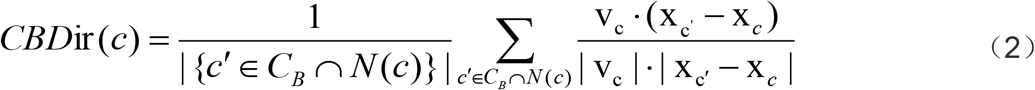

where*Vc, X*_*c*’_ and *X*_*c*_ are vectors representing computed velocity and positions of cell c and c’ in a low-dimensional space. *X* _c’_ − *X* _*c*_ is the cell displacement in this space during the short time.

#### 2. In-cluster coherence (ICCoh) ^17^

ICCoh is computed by assessing the cosine similarity score of the velocity among cells within the same cluster, which evaluates the consistency of the inferred direction of the same cluster regardless of the correctness of the direction. This metric reflects the smoothing of velocities within the cluster. The formula for computing ICCoh score is

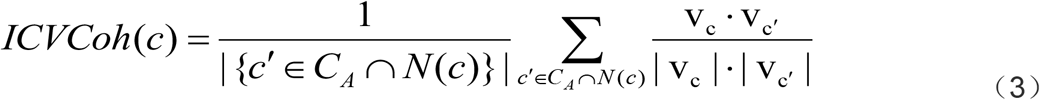

#### 3. Velocity consistency^5^

The score is defined as the average cosine similarity of the velocity vectors to their neighbors. For each cell i,

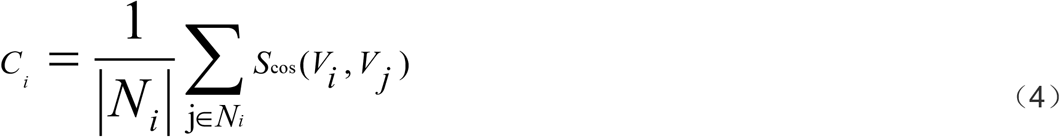

where N_i_ is the 30 nearest neighbor cells of cell i, S_cos_ represents the cosine similarity operation. V_i_, V_j_ are the estimated velocities.

#### 4. Method agreement A1 and A2^25^

This metric is used to assess the consistency of vector predictions for different RNA velocity methods. This metric produces two scores. One is the method agreement score A1, which calculates the cosine similarity of the cell transition vectors created by two different RNA velocity methods. For each pair of methods and each cell, this yields a method agreement score A1.

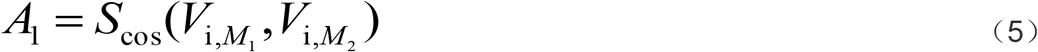

Where S_cos_ represents the cosine similarity operation. V_i, M1_ is the state transition vector of method M1 for cell i and V_i, M2_ is the state transition vector of method M2 for cell i.

The other is method agreement score A2, that is to calculate the cosine similarity between the transition vector of cell i in method M1 and the median vector of cell i. (that is, the cosine similarity is computed between the transition vector of cell i in method M1 and the median vector of cell i.).

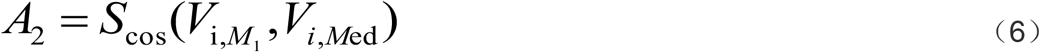

Where V_i, Med_ is the median vector for the cell i. We calculate the central vector for each cell as the median transition vector across all methods. (The median vector of cell i is determined by calculating the median of the transition vectors of cell i across all methods.)

Notably, Dynamo and cellDancer were excluded from this metric assessment due to their lack of cell transfer probability vector calculations.

### Acquisition and preprocessing of datasets

We collected real datasets by searching and selecting 14 existing RNA velocity methods where cells are in dynamic development and differentiation. We incorporated 17 scRNA-seq datasets (Table 1) for evaluation, including datasets used in at least 3 RNA velocity methods, such as pancreatic endocrinogenesis (Dataset1), mouse dentate gyrus (Dataset2), erythrocyte maturation (Dataset3), intestinal organoids (Dataset5), human first post-conception 10th week forebrain (Dataset10), mouse cortical neurons (Dataset13), mouse retinal development (Dataset6), and oligodendrocyte differentiation in the mouse hindbrain (Dataset14). To assess the ability to capture complex transcriptional dynamics, we analyzed a mouse hindbrain dataset with diverse lineage-specific cell populations (Dataset7), a human bone marrow dataset displaying transcriptional enhancements in cell trajectories (Dataset4), a mouse chondrocyte cell development dataset illustrating multiple differentiation pathways (Dataset8), and a human peripheral blood cell dataset representing maturation state (Dataset11). Furthermore, we evaluated the model’s performance on a mouse bone marrow dataset with notably lower UMI counts (Dataset15) to assess its performance with low coverage data. Additionally, we included datasets on developmental and hematopoietic cell dynamics processes: the developing human cortex dataset (Dataset9), the mouse embryonic brain dataset (Dataset16), the human hematopoietic stem cell differentiation dataset (Dataset12), and the mouse hematopoietic stem cell differentiation dataset (Dataset17) to further evaluate the efficacy of different RNA velocity methods.

All scRNA-seq data we used in this study were downloaded publicly(see the “Data availability” section for details on the download sources of the datasets).The original accession numbers and the number of cells and genes in the datasets are described in Table 1.For instance, the mouse dentate gyrus neurogenesis data, we followed the gene and cell filtering methods by Bergen et al. in the scvelo study^5^ and selected 2930 cells with 13913 genes. In the preprocessing phase of RNA velocity analysis, all datasets were log-normalized for the first 3000 highly variable genes, and first and second moments were computed with 30 principal components and 30 nearest neighbors.

### Down-sampling

We used 4 benchmark datasets from 17 datasets (that is, Dataset2: pancreatic endocrinogenesis, Dataset2: dentate gyrus nerve, Dataset3: erythroid maturation, Dataset4: human bone marrow) to test the impact of down-sampling of the splicing matric on each RNA velocity method. Specifically, cells in the spliced matrix of each dataset were sampled stratified by cell cluster (sample rates= 0.4, 0.5, 0.6, 0.7 and 0.8). We down sampled each dataset 5 times at different rates to avoid errors caused by random selection.

### Brief introduction and parameter setting of methods

We evaluated the performance of 14 RNA velocity methods (Table 2). The parameters of each method were set as described below for each program.

### scvelo and velocyto

We followed the guidelines on the scvelo website: https://scvelo.readthedocs.io/en/stable. There are three approaches for estimating RNA velocity in scvelo, the steady-state model (using steady-state residuals), the stochastic model (using second-order moments), and the kinetic model (using a likelihood-based framework). For scvelo-sto and scvelo-dyn, we ran scvelo.tl.velocity with mode=‘stochastic’ and mode=‘dynamic’ with default parameters. The steady state model used in velocyto estimates the velocity. Thus we ran this model with scvelo.tl.velocity mode=‘deterministic’ using default parameters.

### Multivelo

We followed the guidelines on the GitHub repository of Multivelo: https://github.com/welch-lab/MultiVelo. Our dataset is conventional single-cell RNA sequencing data with no ATAC-sequencing (scATAC-seq) data, so we set the parameter rna_only= True.

### veloAE

We followed the guidelines on the veloAE GitHub repository: https://github.com/qiaochen/VeloAE/blob/main/notebooks. We ran the model following the default settings.

### veloVI

We followed the guidelines on the GitHub repository of veloVI: https://github.com/YosefLab/velovi_reproducibility. We ran the model following the default settings.

### veloVAE

We followed the guidelines on the veloVAE GitHub repository: https://github.com/welch-lab/VeloVAE. We set to train a continuous veloVAE model.

### uniTvelo

We followed the guidelines on the GitHub repository of uniTvelo: https://github.com/StatBiomed/UniTVelo/blob/main/notebooks/. we ran the model in “unified” mode by setting velo.FIT_OPTION = ‘1’.

### pyro-velocity

We followed the guidelines on the pyro-velocity GitHub repository: https://github.com/pinellolab/pyrovelocity. We ran the model following the default settings.

### latentvelo

We followed the guidelines on the GitHub repository of latentvelo: https://github.com/Spencerfar/LatentVelo/blob/main/paper_notebooks. We ran the model following the default settings.

### Dynamo

We followed the guidelines on the Dynamo GitHub repository: https://github.com/aristoteleo/dynamo-tutorials. We ran the model following the default settings.

### cell2fate

We followed the guidelines on the GitHub repository of cell2fate: https://github.com/AlexanderAivazidis/cell2fate_notebooks. For datasets with more than 10000 cells, we used the compute_and_plot_total_velocity_scvelo () function to compute velocity.

### cellDancer

We followed the guidelines on the cellDancer GitHub repository: https://github.com/GuangyuWangLab2021/cellDancer. We ran the model following the default settings.

### Deepvelo

We followed the guidelines on the GitHub repository of Deepvelo: https://github.com/bowang-lab/DeepVelo. We ran the model following the default settings.

### Computer platform

We ran CPU tests of the 4 RNA velocity methods (velocyto, scvelo-sto, scvelo-dyn and dynamo) on a computer cluster with Intel(R) Xeon(R) Silver 4210 CPU (2.2 GHz, 14.08 MB L3 cache, 40 CPU cores in total) and 125G memory. The 6 methods, veloAE, veloVI, veloVAE, latentVelo, cellDancer and Deepvelo, are deep learning models that support GPU processing, and the other 4 methods, Multivelo, uniTvelo, pyro-velocity and cell2fate, also support GPU processing. The GPU tests for the 10 methods were performed on a computer with Intel(R) Xeon(R) Gold 6230 CPU (2.1 GHz, 55 MB L3 cache, 80 CPU cores in total), 1TB memory, and NVIDIA A800 GPU (80 GB of memory).

## Data availability

The raw and processed data for all the datasets we use are publicly available. The dentate gyrus neurogenesis data is integrated in scvelo via scv.datasets.dentategyrus() or the original work^31^ under GEO accession number GSE95753. The pancreatic endocrinogenesis data is integrated in scvelo via scv.datasets.pancreas() or the original work^32^ under GEO accession number GSE132188. The erythroid maturation data is integrated in scvelo via scv.datasets.gastrulation_erythroid() or the original work^33^ under GEO accession number GSE87038. The human bone marrow data is integrated in scvelo via scv.datasets.bonemarrow() or the original work^34^ through the Human Cell Atlas data portal under INSDC project accession number ERP120467. The human peripheral blood mononuclear cells data is integrated in scvelo via scv.datasets.pbmc68k() or the original work^35^ under Sequence Read Archive accession code SRP073767.The Intestinal organoid data can be accessed at https://www.dropbox.com/s/25enev458c8egn7/organoid.h5ad? dl=1 or the original work^36^ under GEO accession number GSE128365. The mouse retina development data can be accessed at http://pklab.med.harvard.edu/peterk/review2020/examples/retina/ or the original work^37^ under GEO accession number GSE122466.The mouse hindbrain (GABA, Glial) data can be accessed at https://doi.org/10.6084/m9.figshare.24716592 or the original work^38^ under GEO accession number GSE118068. The mouse organogenesis data can be accessed at https://doi.org/10.6084/m9.figshare.24716592 or the original work^39^ under GEO accession number GSE119945.The mouse cortical neuron data is integrated in dynamo via dyn.sample_data.scNT_seq_neuron_splicing() or the original work^30^ under GEO accession number GSE141851. The human forebrain data can be accessed at https://drive.google.com/file/d/1Bi5Ss7FtyDNV_gZOeoBPittFDf_EHsPw/view or the original work^4^ under Sequence Read Archive accession code SRP129388. The mouse Oligodendrocyte differentiation data can be accessed at http://pklab.med.harvard.edu/ruslan/velocity/oligos/ or the original work^40^ under Sequence Read Archive accession code SRP135960. The mouse bone marrow data can be accessed at https://cell2fate.cog.sanger.ac.uk/browser.html or the original work^7^ under GEO accession number GSE109989. The mouse embryonic brain data can be accessed at https://doi.org/10.6084/10X_multiome_mouse_brain.loom or the original work at https://www.10xgenomics.com/datasets. The human hematopoietic Stem/Progenitor Cell data can be accessed at https://figshare.com/articles/dataset/Human_HSPC_RNA_Data/22575358 or the original work^10^ under GEO accession number GSE209878. The development of human cerebral cortex data can be accessed at https://figshare.com/articles/dataset/Developing_Human_Cortex_RNA_Data/22575376 or the original work^41^ under GEO accession number GSE162170.The mouse hematopoiesis data can be accessed at https://zenodo.org/records/6110279 or the original work^42^ under GEO accession number GSE81682.

## Acknowledgements

This work was supported by the Fundamental Research Funds for the National Natural Science Foundation of China [82272944to QL]

