## Supplementary Figures for "Benchmarking RNA velocity methods across 17 independent studies"

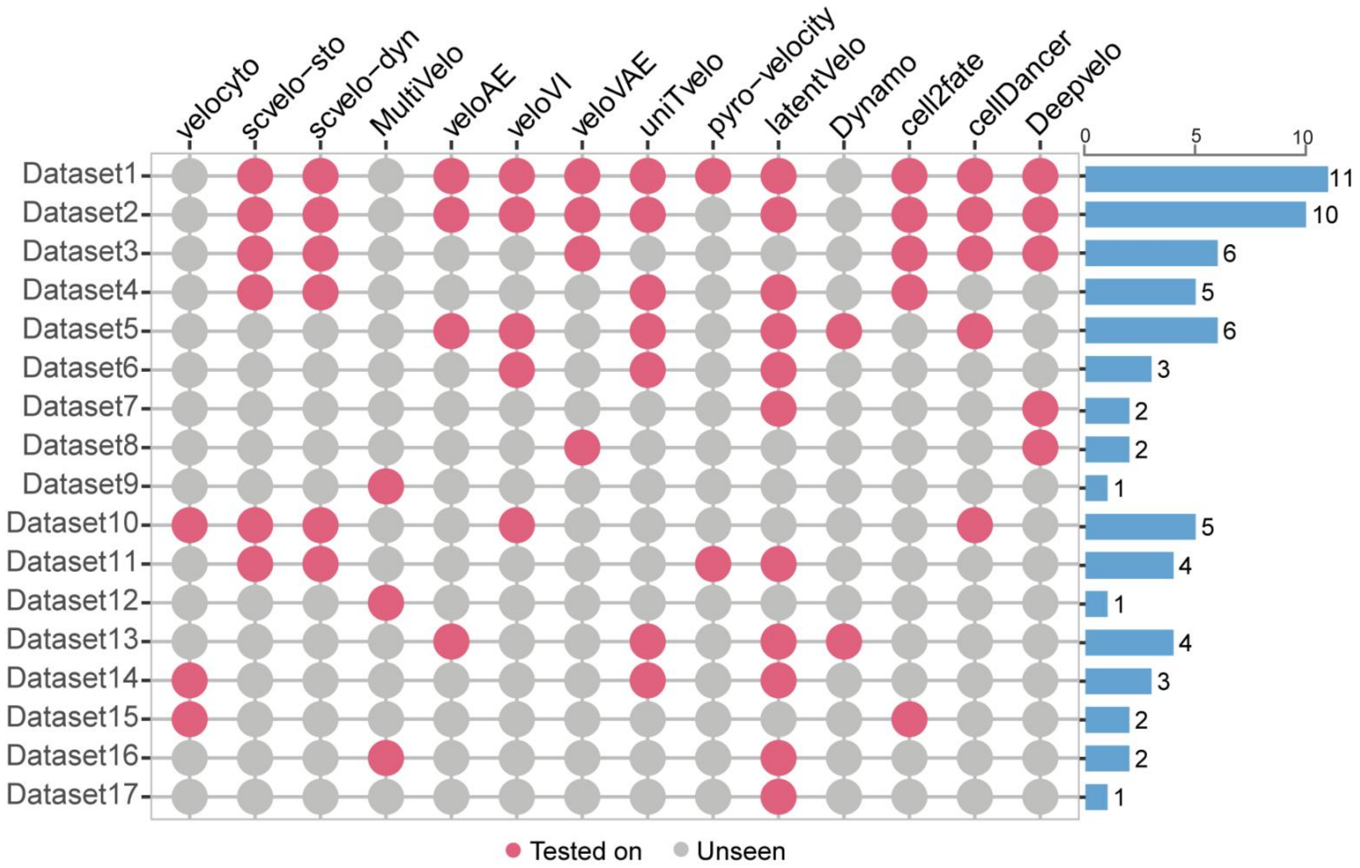


**Supplementary Figure 1. Overview of the usage of 17 datasets across various RNA velocity methods.** The left panel depicts the distribution of dataset usage among different methods, while the bar plot on the right shows the frequency each dataset is used by 14 RNA velocity methods. “Tested on” indicates datasets evaluated in the original publications, whereas “Unseen” denotes datasets not tested by the original authors of each method.

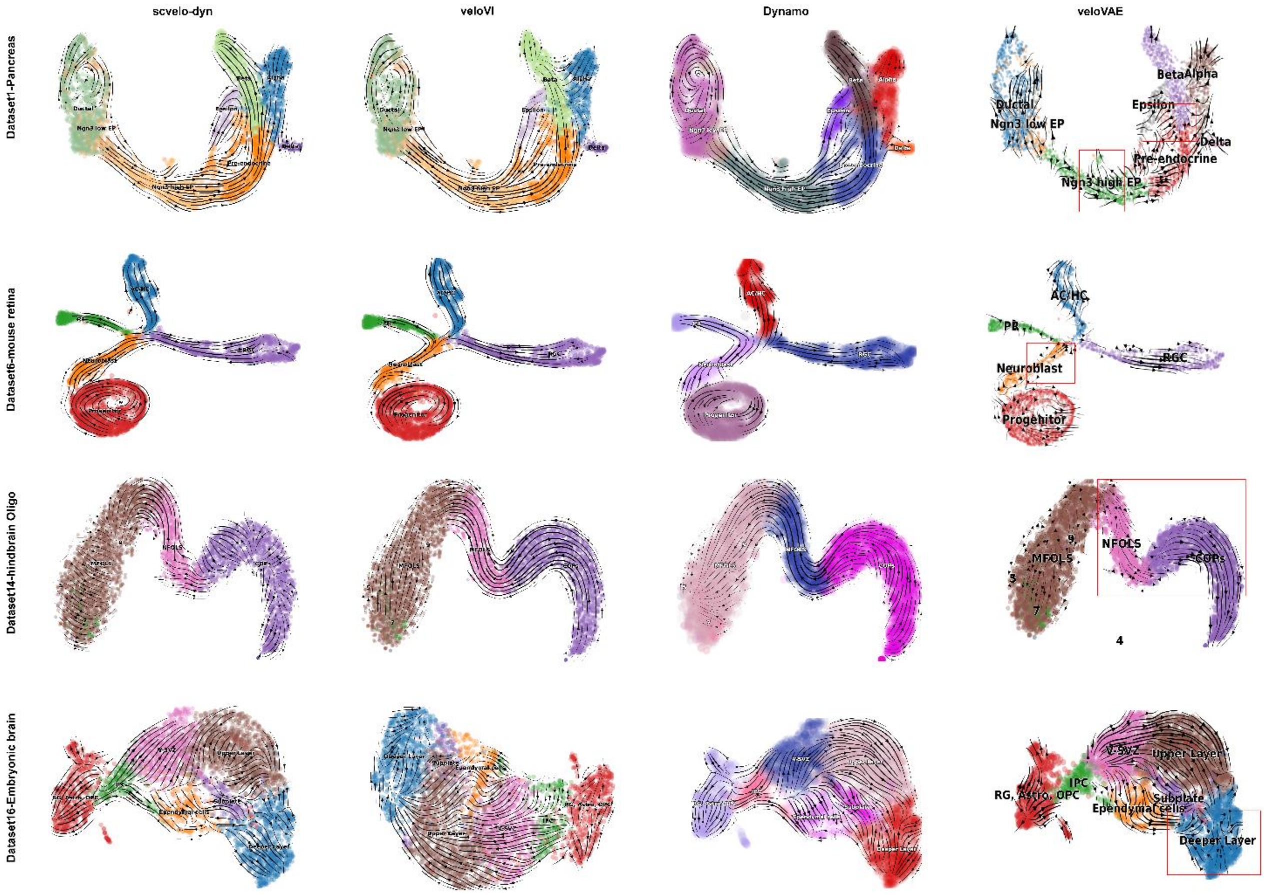


**Supplementary Figure 2. Comparison of velocity field streamlines projected onto UMAP for 4 datasets.** Results from scvelo-dyn, veloVI, Dynamo and veloVAE are shown for dataset1-Pancreas (top), dataset6-Mouse retina (middle I), dataset14-Mouse hindbrain Oligo (middle II) and dataset 16-Embryonic mouse brain (bottom). The regions highlighted by red box indicate reverse cell trajectories inferred by veloVAE.

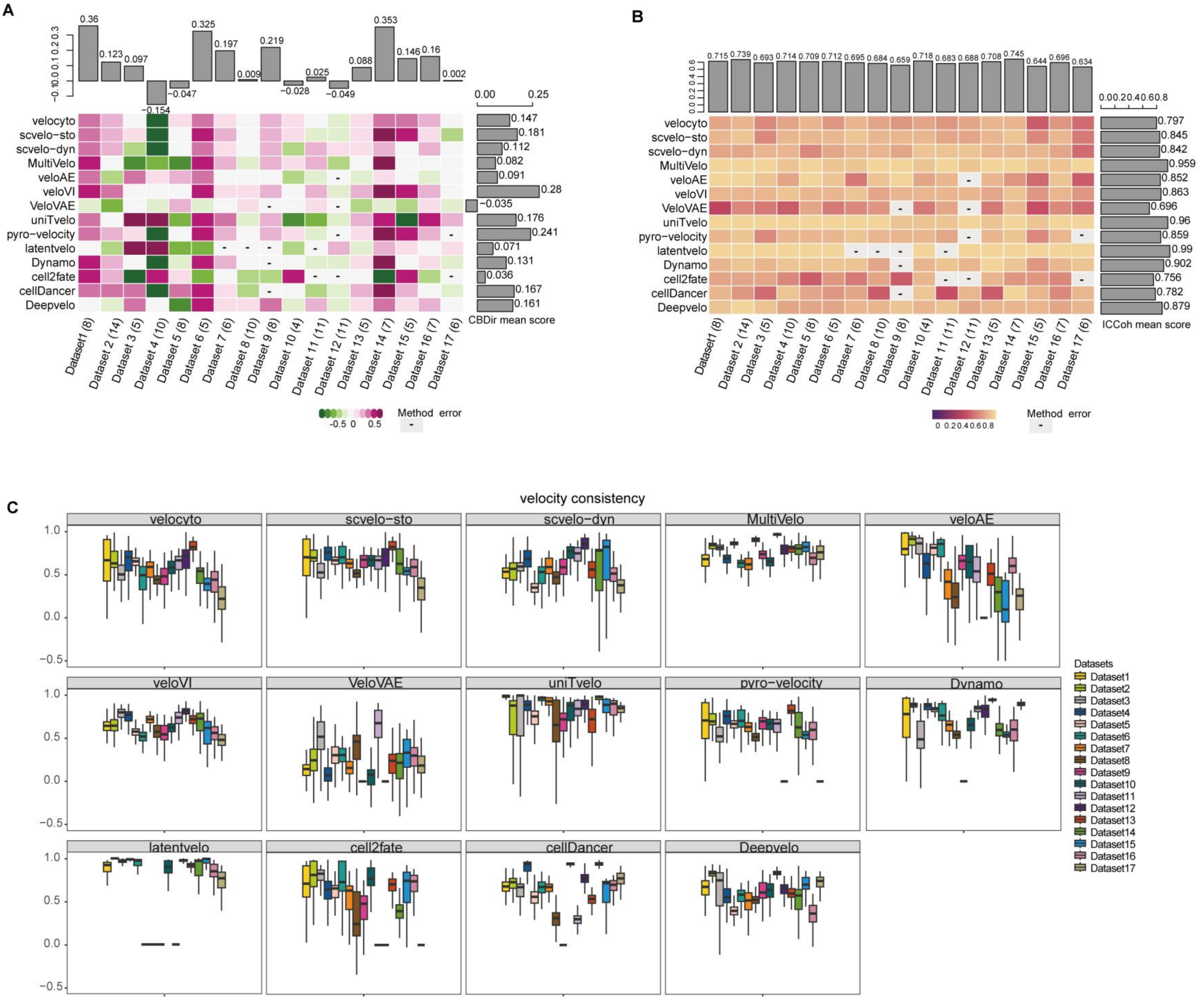


**Supplementary Figure 3. Performance evaluation of 14 RNA velocity methods on 17 scRNA-seq datasets. A.** Comparison of cross-boundary direction correctness (CBDir) scores, histograms of the average CBDir scores across all datasets in each method (right), and histograms of the average CBDir scores across all methods in each dataset (top). **B.** Comparison of intra cluster coherence (ICCoh) scores, histograms of the average ICCoh scores across all datasets in each method (right), and histograms of the average ICCoh scores across all methods in each dataset (top). The number in parentheses in the dataset ID indicates how many cell types are in the dataset. **C.** Boxplot of velocity consistency scores. Colors correspond to different datasets.

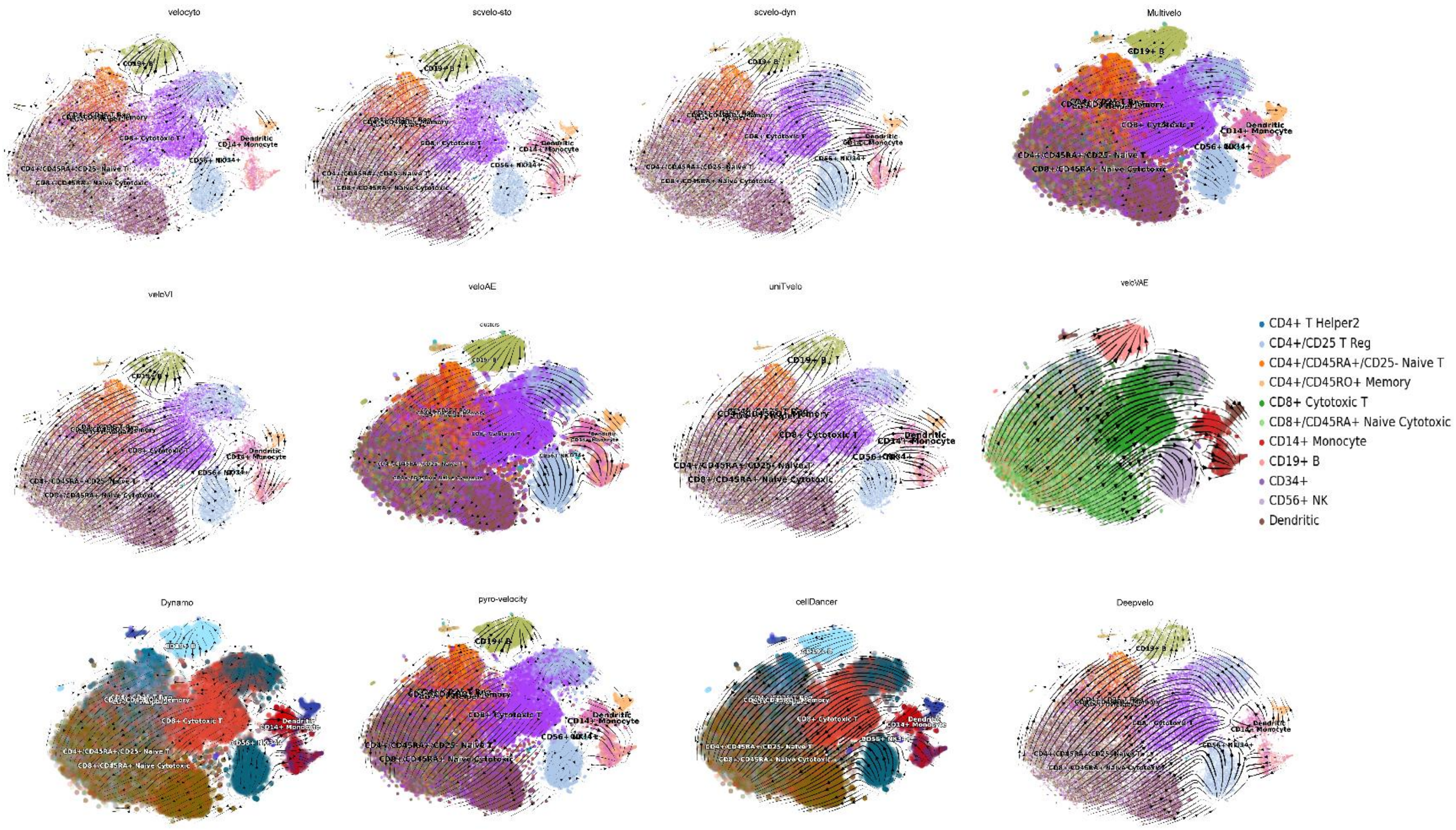


**Supplementary Figure 4. Visualization of velocity field streamlines inferred by different methods on Dataset 11-PBMC 8k, projected onto UMAP embeddings.**

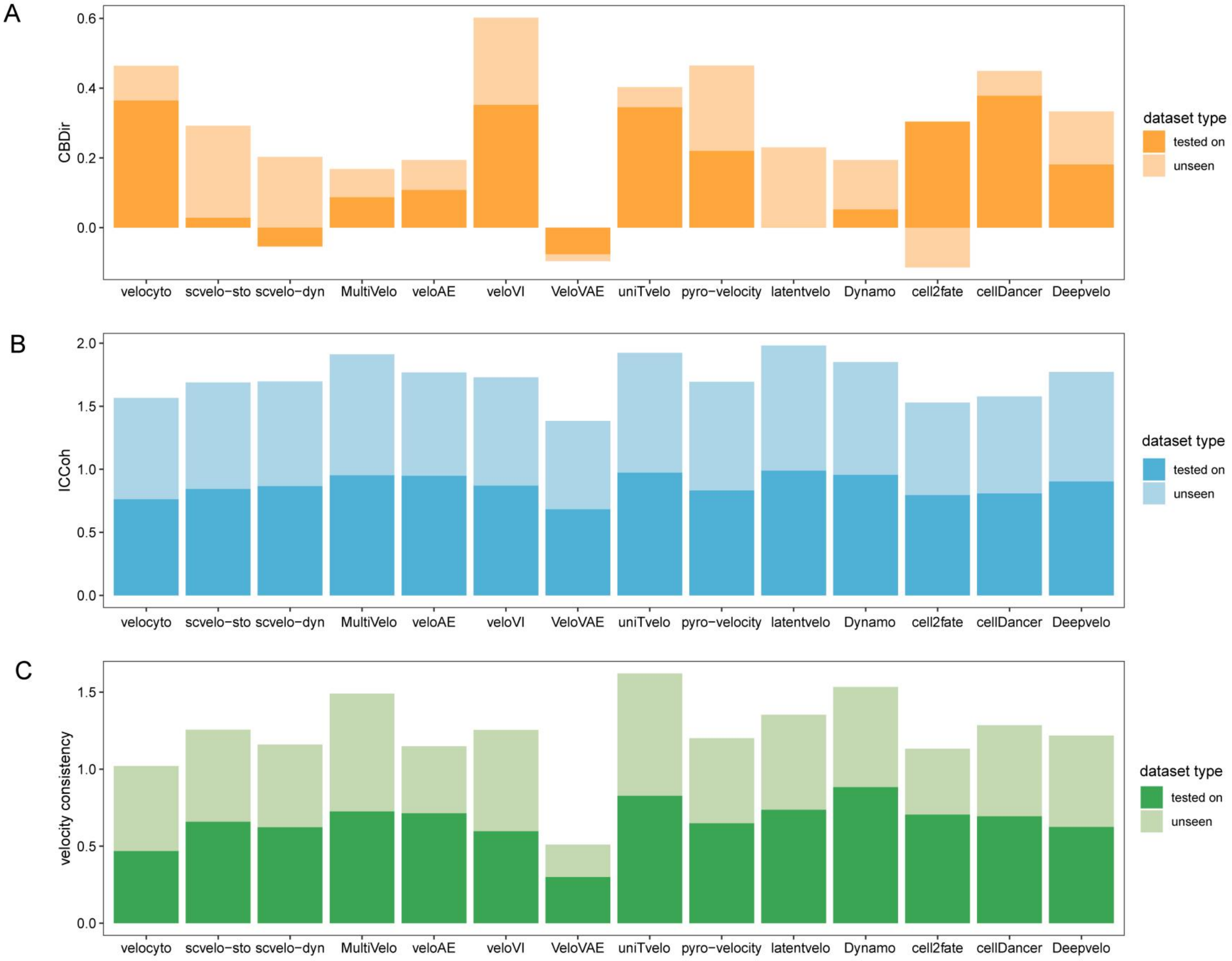


**Supplementary Figure 5. Stacked histograms of performance comparisons for each method by dataset category. A**. CBDir scores. **B**. ICCoh scores. **C**. Velocity consistency scores. “Tested on” and “Unseen” denote the mean values for datasets previously evaluated or not assessed in the original publications.

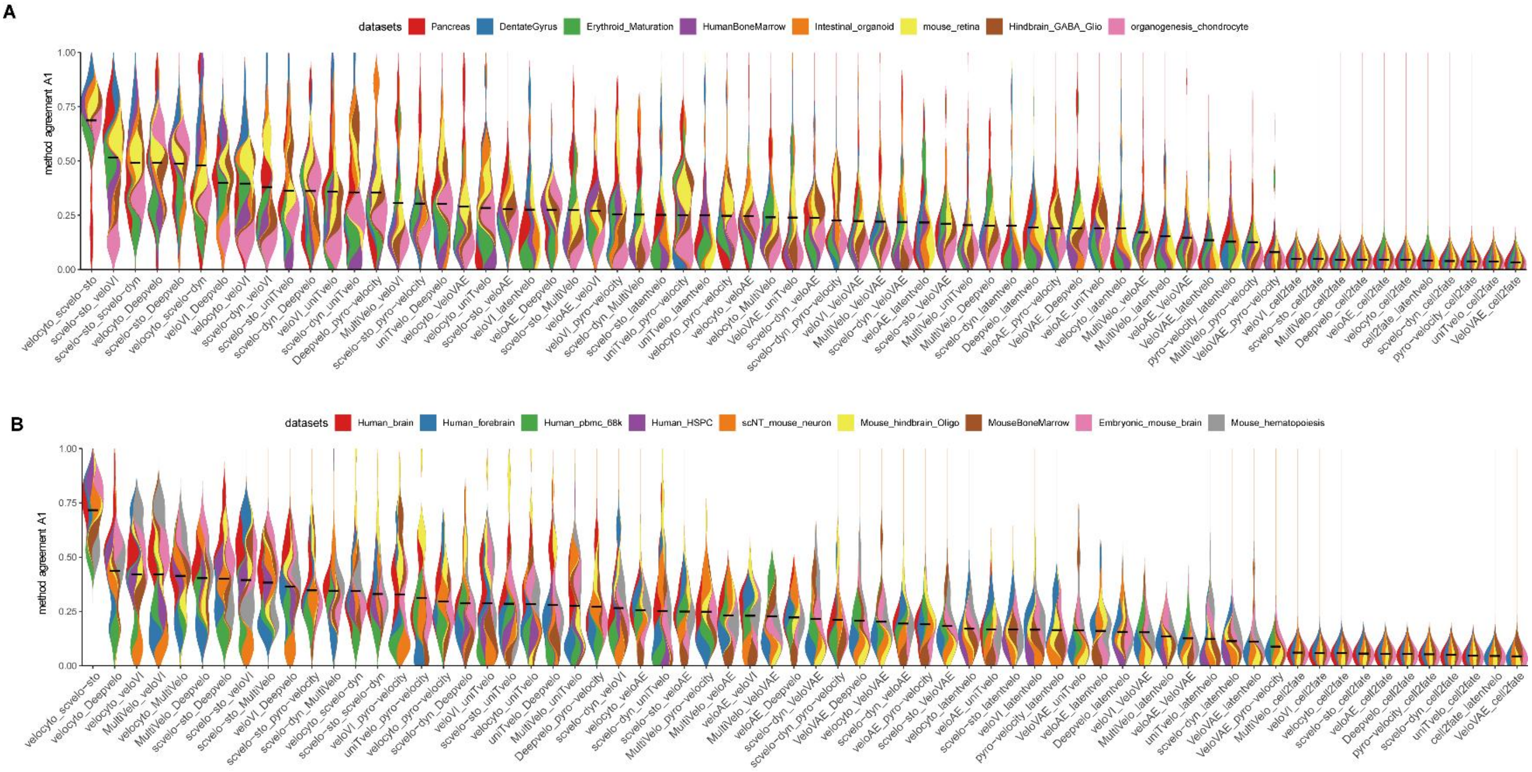


**Supplementary Figure 6. Pairwise comparisons of method agreement A1 scores across 17 datasets. A**. dataset 1-dataset 8. **B**. dataset9- dataset17. All results are color-coded based on datasets, with the black line representing the mean value for each pair of methods.

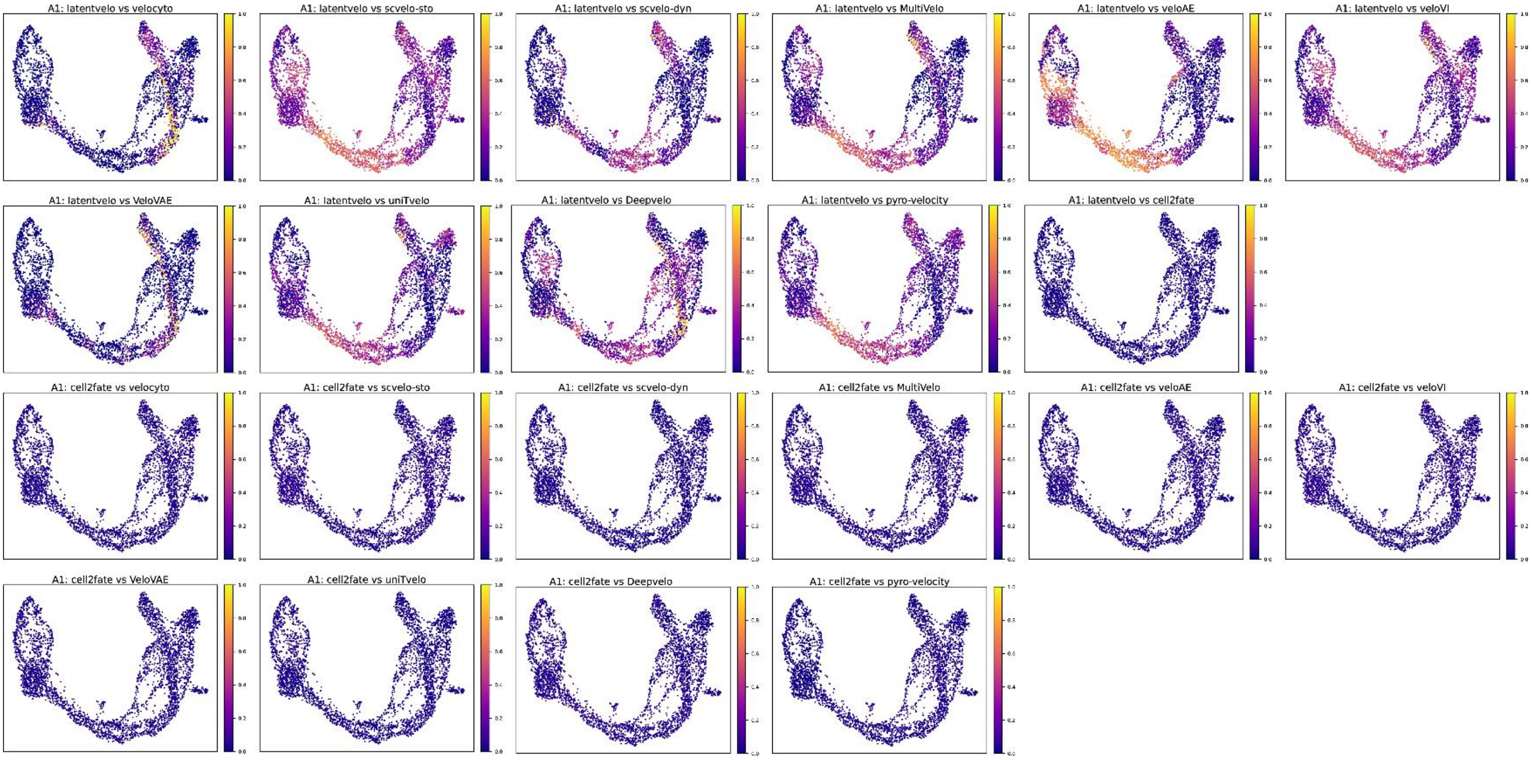


**Supplementary Figure 7. UMAP plots for dataset1, colored by pairwise A1 agreement scores between latentVelo and other methods, as well as between cell2fate and other methods.**

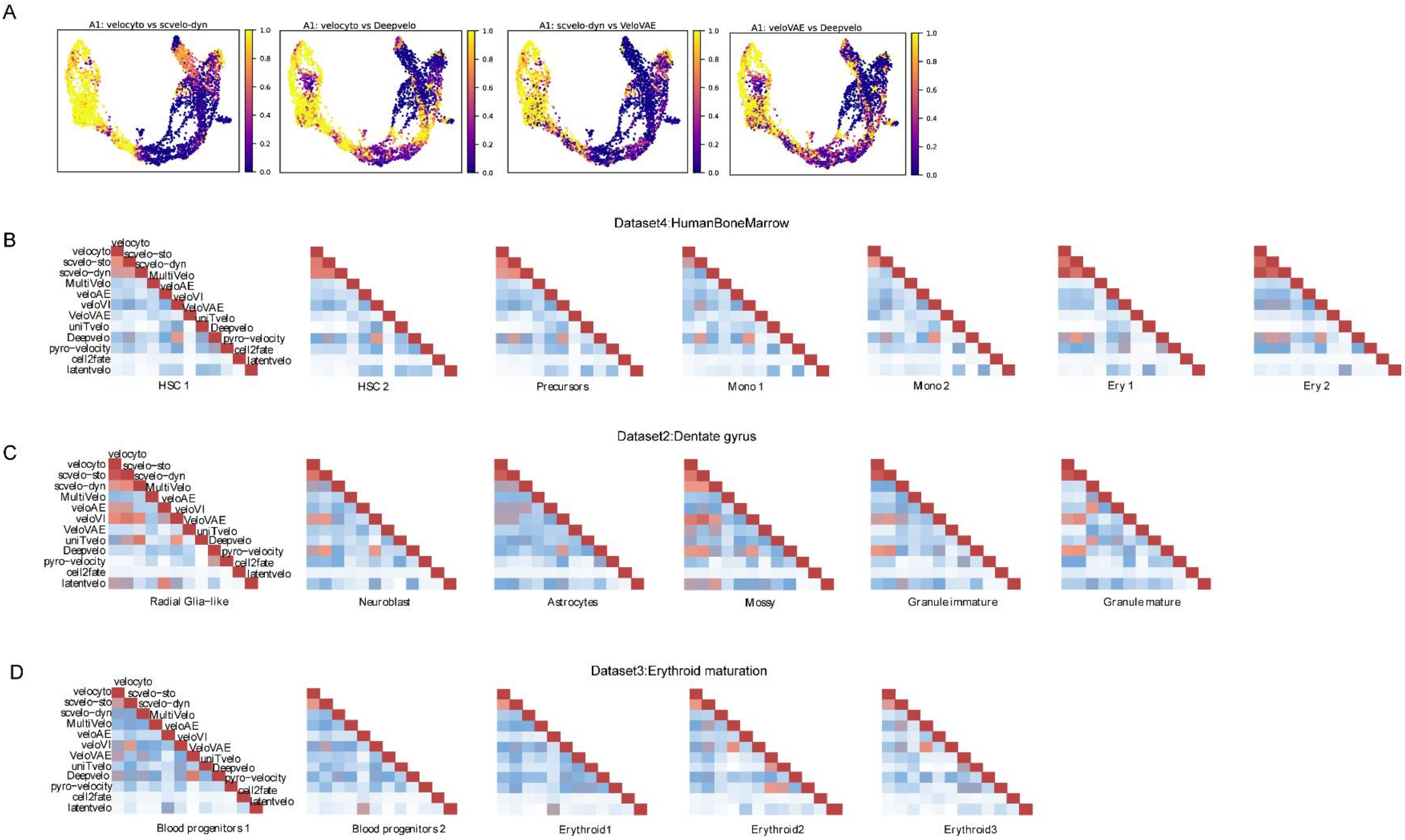


**Supplementary Figure 8. Pairwise comparisons of mean agreement scores (A1) for all methods. A**. UMAP plots for selected paired methods in dataset1 (pancreatic endocrinogenesis), colored by method agreement A1 scores (right). Pairwise comparisons of all methods were performed for the corresponding cell types in **B**. dataset4-human bone marrow. **C**.dataset2-dentate gyrus and **D**. dataset3- erythroid maturation. Heatmaps were colored by the average agreement (A1) of each method within each cell type.

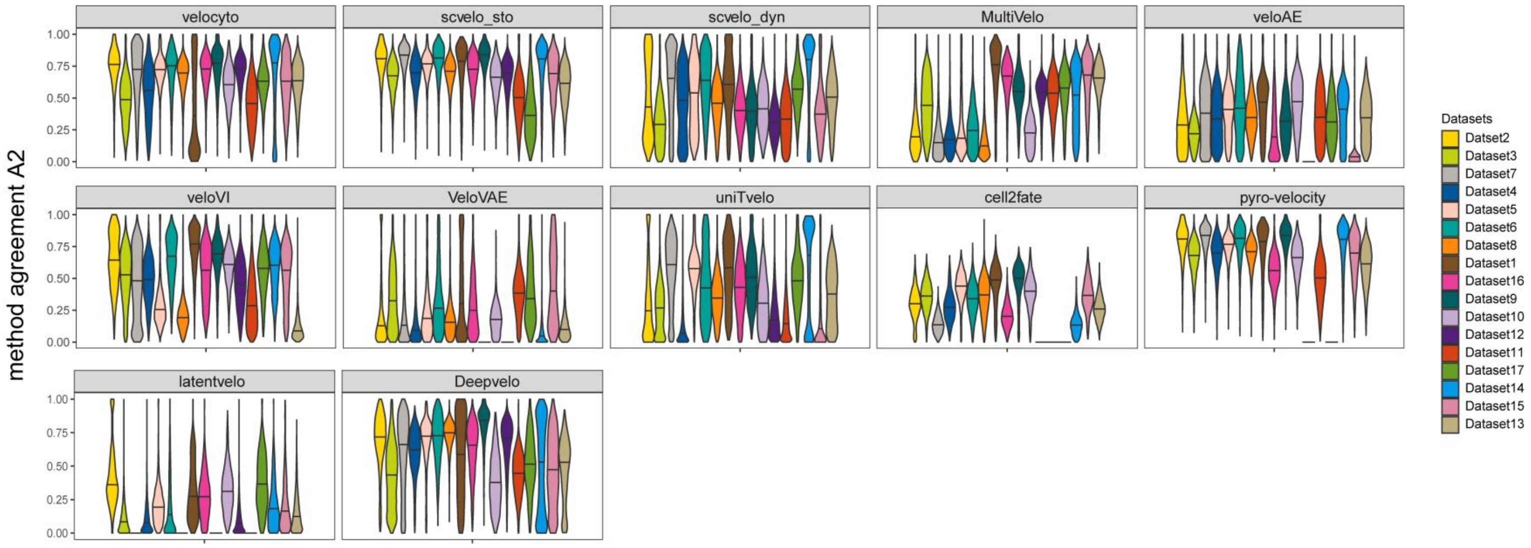


**Supplementary Figure 9. Violin plots of method agreement A2 scores from 12 methods across 17 datasets.**

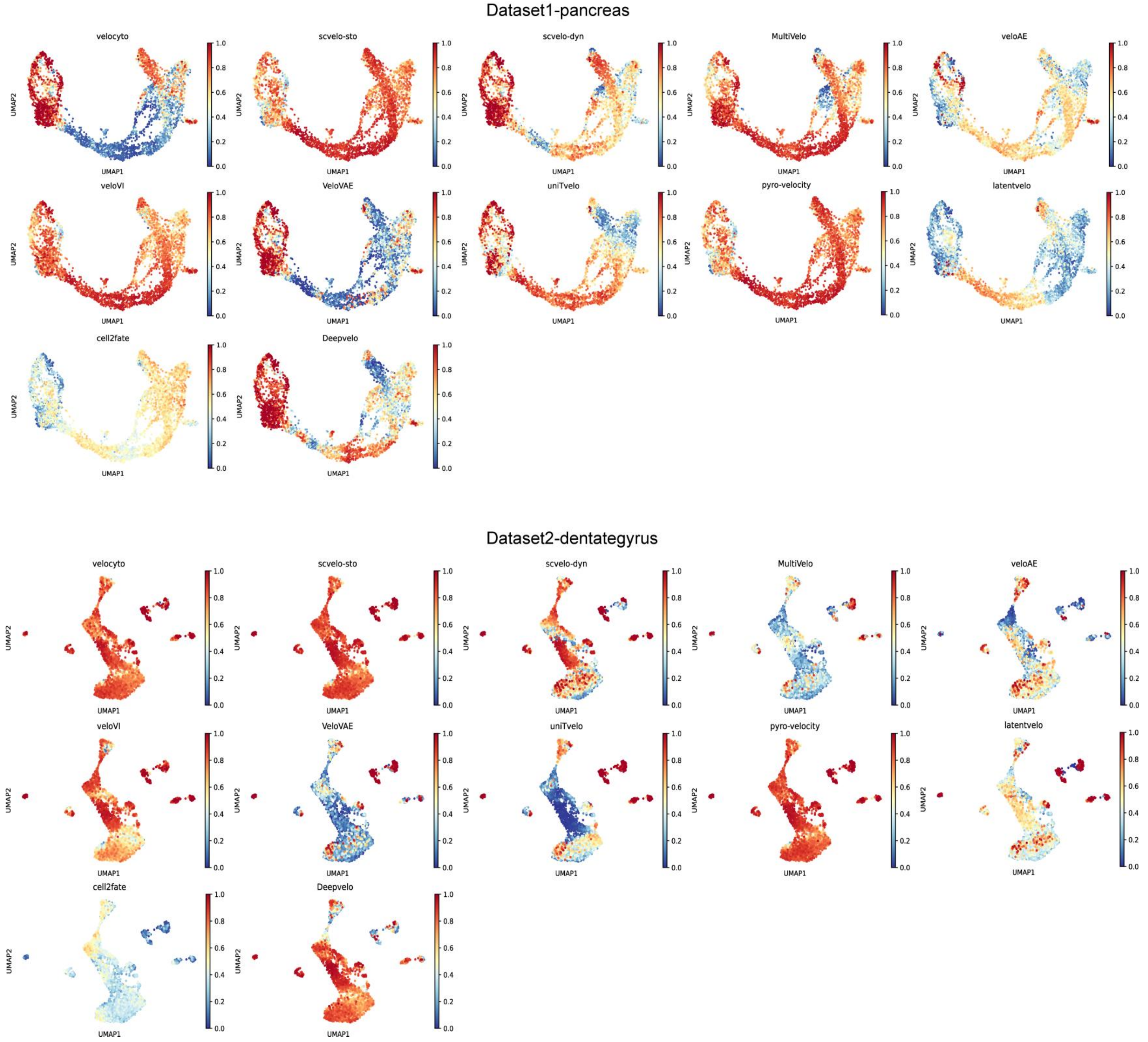


**Supplementary Figure 10. UMAP plots of 12 methods in different datasets. A**. Dataset1 (pancreatic endocrinogenesis) and **B**. Dataset2 (Dentate gyrus), colored by method agreement A2 scores.

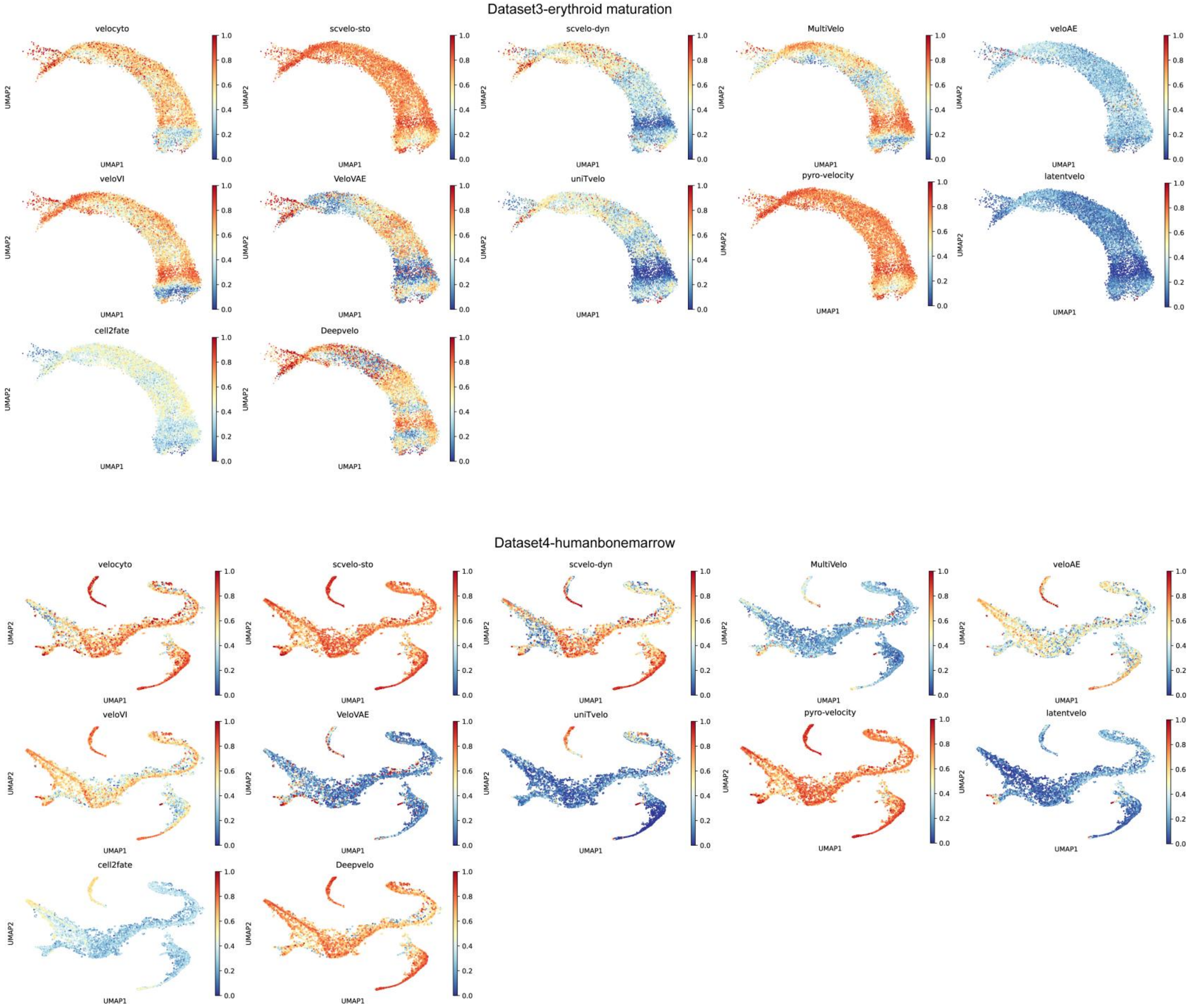


**Supplementary Figure 11. UMAP plots of 12 methods in different datasets. A**. Dataset3 (Erythroid maturation) and **B**. Dataset4 (human bone marrow), colored by method agreement A2 scores.

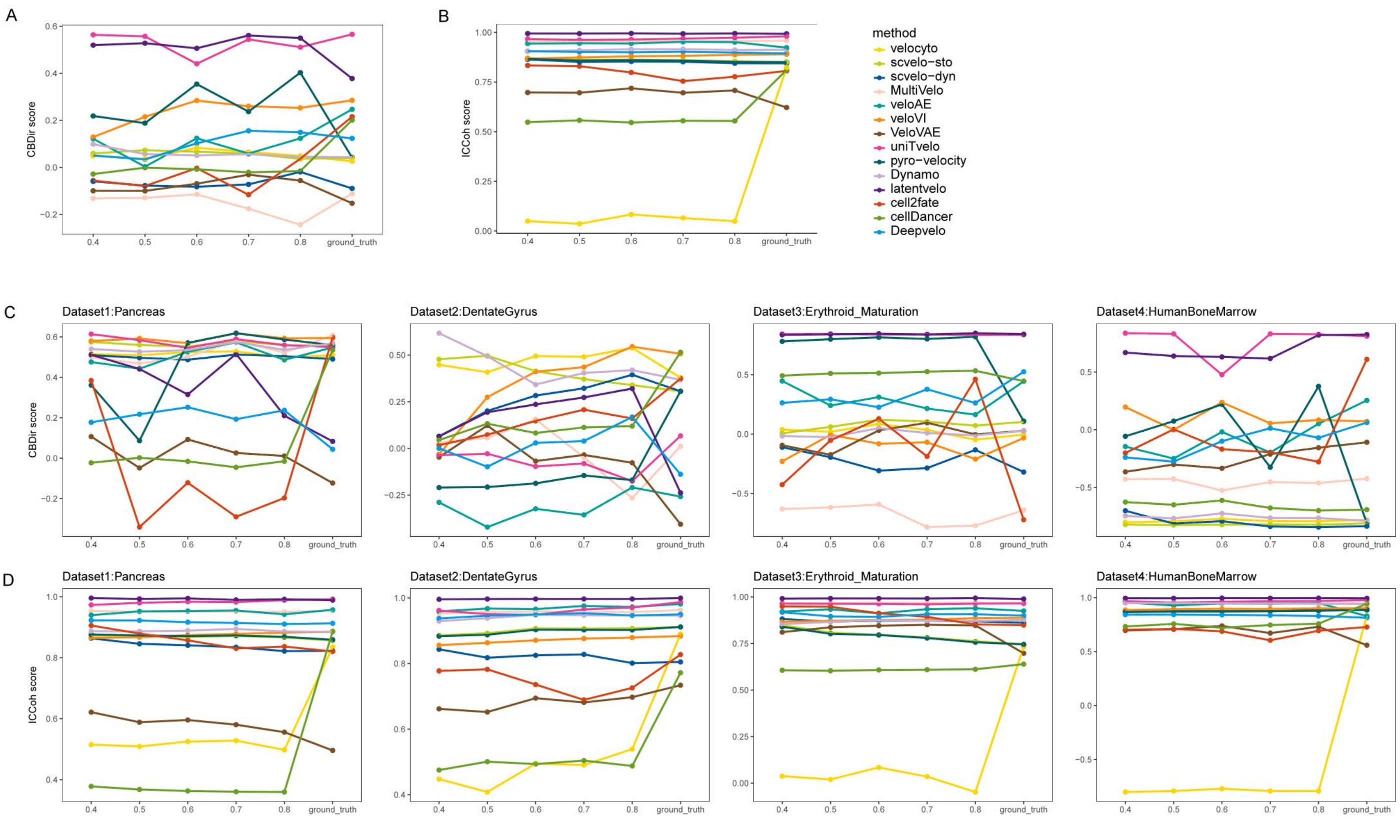


**Supplementary Figure 12. Comparison of CBDir and ICCoh metrics for 14 RNA velocity methods under different sampling rates across 4 benchmark datasets. A-B**. Changes in mean CBDir and ICCoh with sampling ratios; **C-D**. Changes in CBDir and ICCoh for each benchmark dataset at different sampling ratios.

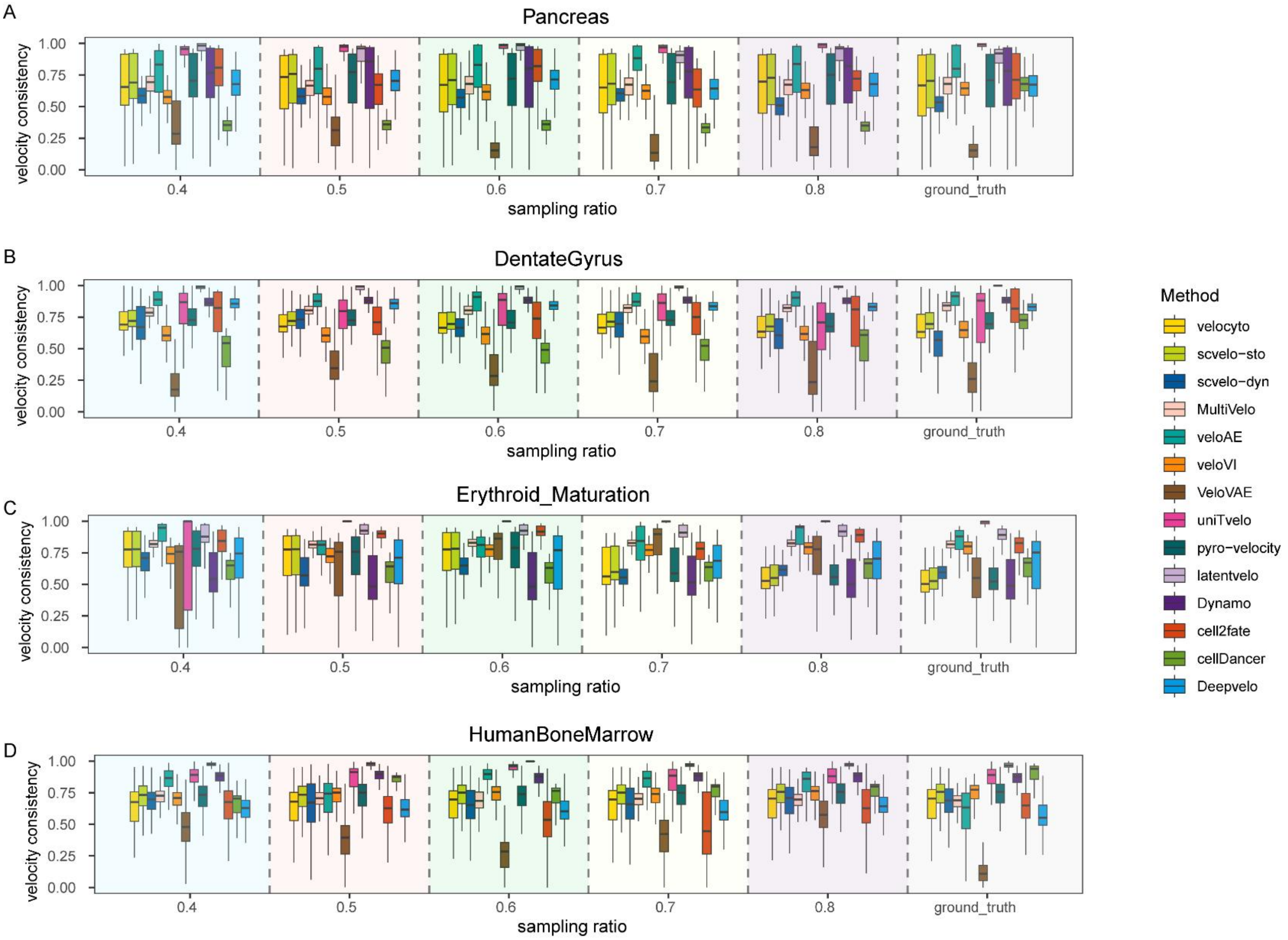


**Supplementary Figure 13. Comparison of velocity consistency metric performance among 14 methods at varying sampling** ratios **across 4 benchmark datasets. A**. Dataset 1 (pancreatic endocrinogenesis). **B**. Dataset 2 (Dentate gyrus). **C**. Dataset 3 (Erythroid maturation). **D**. Dataset 4 (human bone marrow).

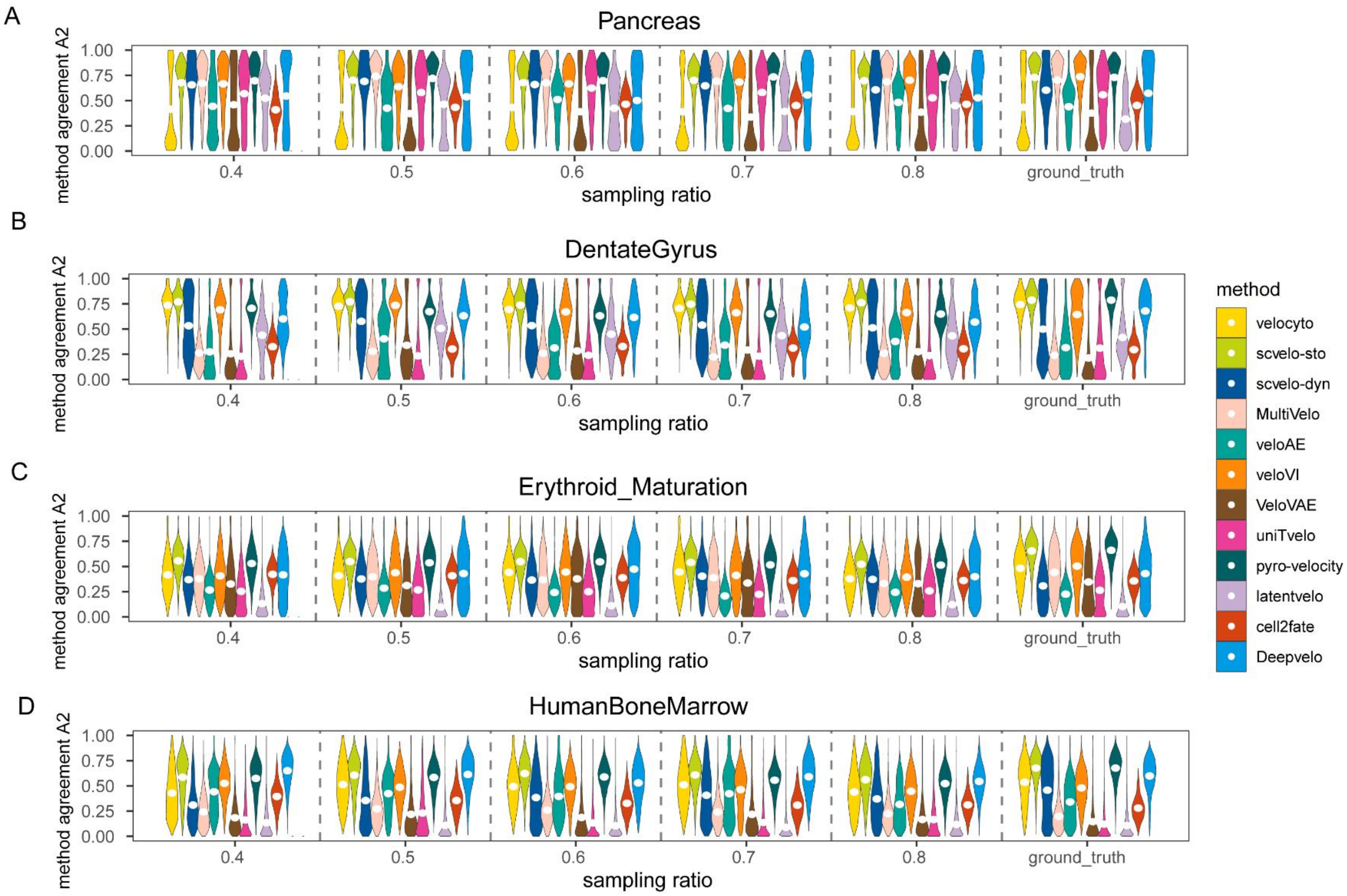


**Supplementary Figure 14. Comparison of method agreement A2 metric performance among 14 methods at varying sampling** ratios **across 4 benchmark datasets. A**. Dataset 1 (pancreatic endocrinogenesis). **B**. Dataset 2 (Dentate gyrus). **C**. Dataset 3 (Erythroid maturation). **D**. dataset4 (human bone marrow).
